# Reconciling functional differences in populations of neurons recorded with two-photon imaging and electrophysiology

**DOI:** 10.1101/2020.08.10.244723

**Authors:** Joshua H. Siegle, Peter Ledochowitsch, Xiaoxuan Jia, Daniel Millman, Gabriel K. Ocker, Shiella Caldejon, Linzy Casal, Andrew Cho, Daniel J. Denman, Séverine Durand, Peter A. Groblewski, Greggory Heller, India Kato, Sara Kivikas, Jerome Lecoq, Chelsea Nayan, Kiet Ngo, Philip R. Nicovich, Kat R. North, Tamina K. Ramirez, Jackie Swapp, Xana Waughman, Ali Williford, Shawn R. Olsen, Christof Koch, Michael A. Buice, Saskia E. J. de Vries

## Abstract

Extracellular electrophysiology and two-photon calcium imaging are widely used methods for measuring physiological activity with single-cell resolution across large populations of neurons in the brain. While these two modalities have distinct advantages and disadvantages, neither provides complete, unbiased information about the underlying neural population. Here, we compare evoked responses in visual cortex recorded in awake mice under highly standardized conditions using either imaging or electrophysiology. Across all stimulus conditions tested, we observe a larger fraction of responsive neurons in electrophysiology and higher stimulus selectivity in calcium imaging. This work explores which data transformations are most useful for explaining these modality-specific discrepancies. We show that the higher selectivity in imaging can be partially reconciled by applying a spikes-to-calcium forward model to the electrophysiology data. However, the forward model could not reconcile differences in responsiveness without sub-selecting neurons based on event rate or level of signal contamination. This suggests that differences in responsiveness more likely reflect neuronal sampling bias or cluster-merging artifacts during spike sorting of electrophysiological recordings, rather than flaws in event detection from fluorescence time series. This work establishes the dominant impacts of the two modalities’ respective biases on a set of functional metrics that are fundamental for characterizing sensory-evoked responses.

## Introduction

Systems neuroscience aims to explain how complex adaptive behaviors can arise from the interactions of many individual neurons. As a result, population recordings—which capture the activity of multiple neurons simultaneously—have become the foundational method for progress in this domain. Extracellular electrophysiology and calcium-dependent two-photon optical physiology are by far the most prevalent population recording techniques, due to their single-neuron resolution, ease of use, and scalability. Recent advances have made it possible to record simultaneously from thousands of neurons with electrophysiology (Jun et al., 2017; Siegle et al., 2019; Stringer et al., 2019a) or tens of thousands of neurons with calcium imaging (Sofroniew et al., 2016; Stringer et al., 2019b; Weisenburger et al., 2019). While insights gained from both methods have been invaluable to the field, it is clear that neither technique provides a completely faithful picture of the underlying neural activity. In this study, our goal is to better understand the inherent biases of each recording modality, and specifically how to appropriately compare results obtained with one method to the other.

Head-to-head comparisons of electrophysiology and imaging data are rare in the literature, but are critically important as the practical aspects of each method affect their suitability for different experimental questions. Since the expression of calcium indicators can be restricted to genetically defined cell types, imaging can easily target recordings to specific sub-populations (Madisen et al., 2015). Similarly, the use of retro- or anterograde viral transfections to drive indicator expression allows imaging to target sub-populations defined by their projection patterns (Glickfeld et al., 2013; Gradinaru et al., 2010). The ability to identify genetically or projection-defined cell populations in electrophysiology experiments is far more limited (Economo et al., 2018; Jia et al., 2019; Lima et al., 2009). Both techniques have been adapted for chronic recordings, but imaging offers the ability to reliably return to the same neurons over many days without the need to implant bulky hardware (Peters et al., 2014). Furthermore, because imaging captures structural, in addition to functional, data, individual neurons can be precisely registered to tissue volumes from electron microscopy (Bock et al., 2011; Lee et al., 2016), in vitro brain slices (Ko et al., 2011), and potentially other *ex vivo* techniques such as *in situ* RNA profiling (Chen et al., 2015). In contrast, the sources of extracellular spike waveforms are very difficult to localize with sufficient precision to enable direct cross-modal registration.

Inherent differences in the spatial sampling properties of electrophysiology and imaging are widely recognized, and influence what information can be gained from each method (Figure 1A). Multi-photon imaging typically yields data in a single plane tangential to the cortical surface, and is limited to depths of <1 mm due to a combination of light scattering and absorption in tissue. While multi-plane (Yang et al., 2016) and deep structure (Ouzounov et al., 2017) imaging are both areas of active research, imaging of most subcortical structures requires physical destruction of more superficial tissues (Dombeck et al., 2010; Feinberg and Meister, 2015; Skocek et al., 2018). Extracellular electrophysiology, on the other hand, utilizes micro-electrodes embedded in the tissue, and thus dense recordings are easiest to perform along a straight line, normal to the cortical surface, in order to minimize per-channel tissue displacement. Linear probes provide simultaneous access to neurons in both cortex and subcortical structures, but make it difficult to sample many neurons from the same cortical layer.

**Figure 1:**
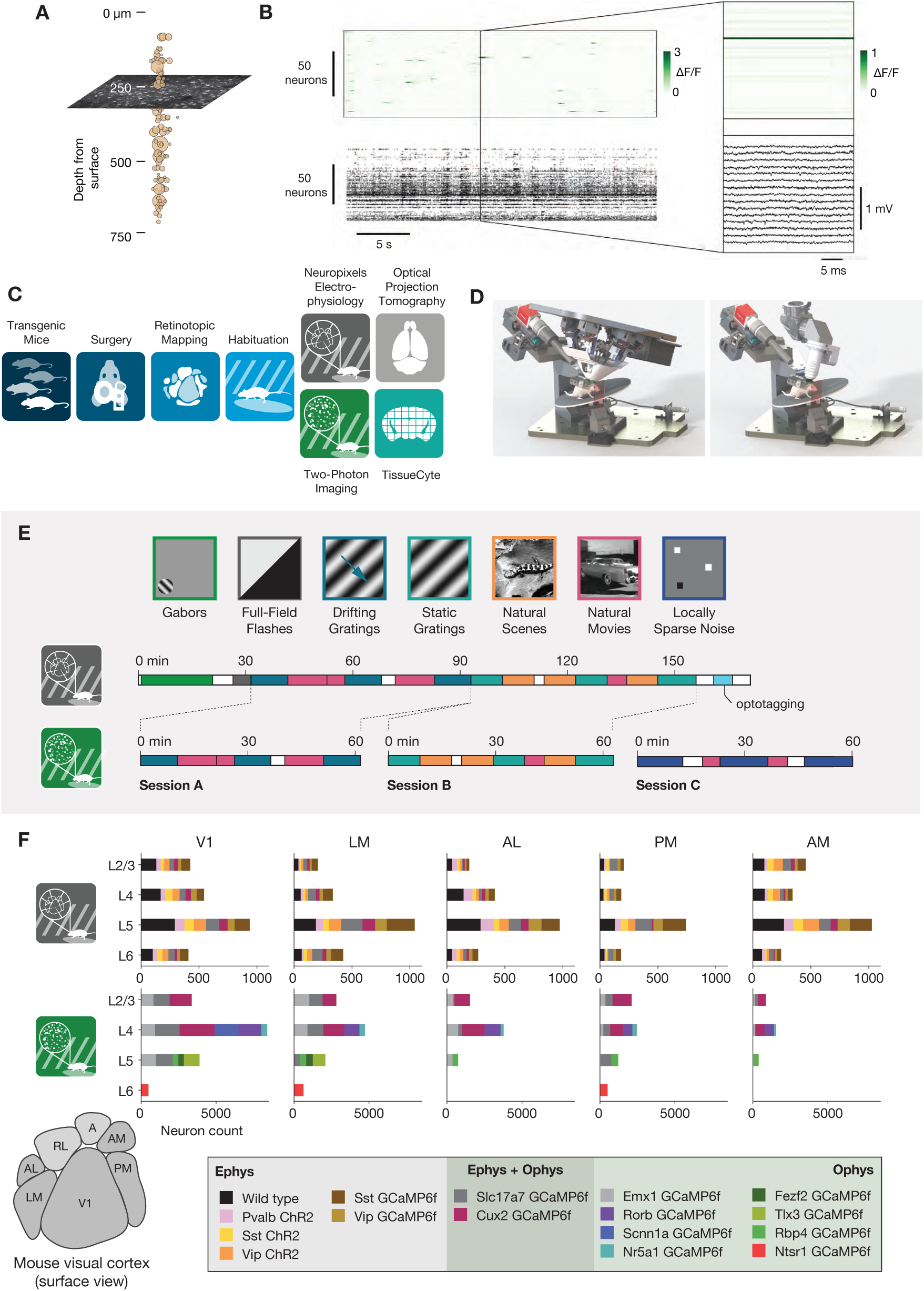
Overview of the ephys and ophys datasets. (A) Illustration of the orthogonal spatial sampling profiles of the two modalities. Black and white area represents a typical ophys imaging plane (at approximately 250 µm below the brain surface), while tan circles represent the inferred locations of cortical neurons recorded with a Neuropixels probe (area is proportional to overall amplitude). (B) Comparison of temporal dynamics between the modalities. *Top:* a heatmap of ΔF/F values for 100 neurons simultaneously imaged in V1 during the presentation of a 30 s movie clip. *Bottom:* raster plot for 100 neurons simultaneously recorded with a Neuropixels probe in V1 in a different mouse viewing the same movie. *Inset:* Close-up of one sample of the ophys heatmap, plotted on the same timescale as 990 samples from 15 electrodes recorded during the equivalent interval from the ephys experiment. (C) Steps in the two data generation pipelines. Following habituation, mice proceed to either two-photon imaging or Neuropixels electrophysiology. (D) Side-by-side comparison of the rigs used for ephys (left) and ophys (right). (E) Schematic of the stimulus set used for both modalities. The ephys stimuli are shown continuously for a single session, while the ophys stimuli are shown over the course of 3 separate sessions. (F) Histogram of neurons recorded in each area and layer, grouped by mouse genotype.

The temporal resolutions of these two methodologies also differ in critical ways (Figure 1B). Imaging is limited by the dwell time required to capture enough photons to distinguish physiological changes in fluorescence from noise (Svoboda and Yasuda, 2006), and the kinetics of calcium-dependent indicators additionally constrain the ability to temporally localize neural activity (Chen et al., 2013). While kilohertz-scale imaging has been achieved (Kazemipour et al., 2019; Zhang et al., 2019), most studies are based on data sampled at frame rates between 1 and 30 Hz. In contrast, extracellular electrophysiology requires sampling rates of 20 kHz or higher, in order to capture the action potential waveform shape that is essential for accurate spike sorting. High sampling rates allow extracellular electrophysiology to pin-point neural activity in time with sub-millisecond resolution, enabling analyses of fine-timescale synchronization across simultaneously recorded neural populations. The fact that electrophysiology can measure action potentials—what we believe to be the fundamental currency of neuronal communication and causation—bestows upon it a more basic ontological status than on calcium imaging, which captures the indirect consequences of a neuron’s spike train.

To date, there has been no comprehensive attempt to characterize how the choice of recording modality affects the inferred functional properties of neurons in sensory cortex. Our limited understanding of how scientific conclusions may be skewed by the recording modality represents the weakest link in the chain of information integration across the techniques available to neurophysiologists today. To address this, we took advantage of two recently collected large-scale datasets that sampled neural activity in mouse visual cortex using either two-photon calcium imaging (de Vries et al., 2020) or dense extracellular electrophysiology (Siegle et al., 2019). These datasets were collected using standardized pipelines, such that the surgical methods, experimental steps, and physical geometry of the recording rigs were matched as closely as possible (Figure 1C,D). The overall similarity of these Allen Brain Observatory pipelines eliminates many of the potential confounding factors that arise when comparing results from imaging and electrophysiology experiments. We note that this is not an attempt at calibration against ground truth data, but rather an attempt to reconcile results across two uniquely comprehensive datasets collected under maximally identical conditions.

Our comparison focused on metrics that capture three fundamental features of neural responses to environmental stimuli: (1) responsiveness, (2) preference (i.e., condition that maximizes the peak response), and (3) selectivity (i.e., sharpness of tuning). Responsiveness metrics characterize whether or not a particular stimulus type (e.g., drifting gratings) reproducibly elicits increased activity. For responsive neurons, preference metrics (e.g. preferred temporal frequency) determine which stimulus condition (out of a finite set) elicits the largest response, and serve as an indicator of a neuron’s functional specialization—for example, whether it responds preferentially to slow- or fast-moving stimuli. Lastly, selectivity metrics (e.g. orientation selectivity, lifetime sparseness) characterize a neuron’s ability to distinguish between particular exemplars within a stimulus class. All three of these features must be measured accurately in order to understand how stimuli are represented by individual neurons.

We find that preference metrics are largely invariant across modalities. However, electrophysiology suggests that neurons show a higher degree of responsiveness, while imaging suggests that responsive neurons show a higher degree of selectivity. In the absence of steps taken to mitigate these differences, the two modalities will yield mutually incompatible conclusions about the basic response properties of the areas under study. These differences can be partially reconciled through a combination of applying a spikes-to-calcium forward model to the electrophysiology data (Deneux et al., 2016) and by sub-selection of neurons based either on event rate or by contamination level (the likelihood that signal from other neurons is misattributed to the neurons under consideration). This reconciliation reveals the respective biases of these two recording modalities, in which extracellular electrophysiology predominantly captures the activity of highly active units while missing low-firing-rate units, while calcium-indicator binding dynamics sparsify neural responses and supralinearly amplify spike bursts.

## Results

We compared the visual responses measured in the Allen Brain Observatory Visual Coding (“ophys”) and Allen Brain Observatory Neuropixels (“ephys”) datasets, publicly available through brain-map.org and the AllenSDK Python package. These datasets consist of recordings from neurons in six cortical visual areas (as well as subcortical areas in the Neuropixels dataset) in the awake, head-fixed mouse in response to a battery of passively viewed visual stimuli. For both datasets, the same drifting gratings, static gratings, natural scenes, and natural movie stimuli were shown (Figure 1E). These stimuli were presented in a single 3-hour recording session for the ephys dataset. For the ophys dataset, these stimuli were divided across three separate 1-hour imaging sessions from the same group of neurons. In both ephys and ophys experiments, mice were free to run on a rotating disc, the motion of which was continuously recorded.

The ophys dataset was collected using genetically encoded GCaMP6f (Chen et al., 2013) under the control of specific Cre driver lines. These Cre drivers limit the calcium indicator expression to specific neuronal populations, including different excitatory and inhibitory populations found in specific cortical layers (see de Vries et al., 2020 for details). The ephys dataset also made use of transgenic mice in addition to wild-type mice. These transgenic mice expressed either channelrhodopsin in specific inhibitory populations for identification using optotagging (see Siegle et al., 2019 for details), or GCaMP6f in specific excitatory or inhibitory populations (see Methods). Unlike in the ophys dataset, however, these transgenic tools did not determine which neurons were recorded.

We limited our comparative analysis to putative excitatory neurons from 5 cortical visual areas (V1, LM, AL, PM, and AM). For ophys, we only included data from 10 excitatory Cre lines, while for ephys we limited our analysis to regular-spiking units by setting a threshold on the waveform duration (>0.4 ms). After this filtering step, we were left with 41,578 neurons from 170 mice in ophys, and 11,030 neurons from 52 mice in ephys. The total number of cells for each genotype, layer, and area is shown in Figure 1F.

### Baseline metric comparison

Prior to computing response metrics for the ophys dataset, events were extracted from the continuous ΔF/F trace for each ROI using an *ℓ*_0_-regularized algorithm (Jewell et al., 2018; Jewell and Witten, 2018; de Vries et al., 2020). This algorithm identified the onset time and the corresponding magnitude of fluorescence change (magnitudes are reported on the same scale as ΔF/F and are therefore unitless). For the most part, such “events” do not represent individual spikes (Ledochowitsch et al., 2019; Huang et al., 2019), but rather are heavily biased towards indicating short bouts of high firing rate, e.g. bursting. Responses were computed using these event magnitudes for the ophys data. For the ephys dataset, responses were computed using the spike times identified by the spike-sorting algorithm, Kilosort2 (Pachitariu et al., 2016; Stringer et al., 2019a).

A comparison of ephys spikes and ophys events shows stark differences between the features of these two modalities. The event rate histograms both display a roughly log-normal distribution (Buzsáki and Mizuseki, 2014), but the most active neurons from ophys rarely have an event rate of more than 1 Hz, which is on the low end of spike rates observed in ephys (Figure 2A). This discrepancy can partially be accounted for by the lower sampling rate for ophys, which limits the maximum event rate. However, this comparison is also confounded by the fact that one event is not equivalent to one spike. There is rich information contained in the event amplitudes, which have a non-linear—albeit on average monotonic—relationship with the underlying number of spikes within a window (Ledochowitsch et al., 2019). Integrating over all 250 ms windows in the ophys dataset (*N* = 40 million intervals with at least one event, out of 1.1 billion possible intervals), event amplitudes vary over 4 orders of magnitude, with most amplitudes falling in the middle of this range (Figure 2B; the truncated shoulder at the lowest end of this range is due to the noise-dependent event detection regularization, see Methods for details). Within the same 250 ms window, the number of ephys spikes follows a much different distribution, with the vast majority of non-silent intervals containing only one or two spikes (Figure 2C; *N* = 129 million intervals with at least one spike, out of 417 million possible intervals).

**Figure 2:**
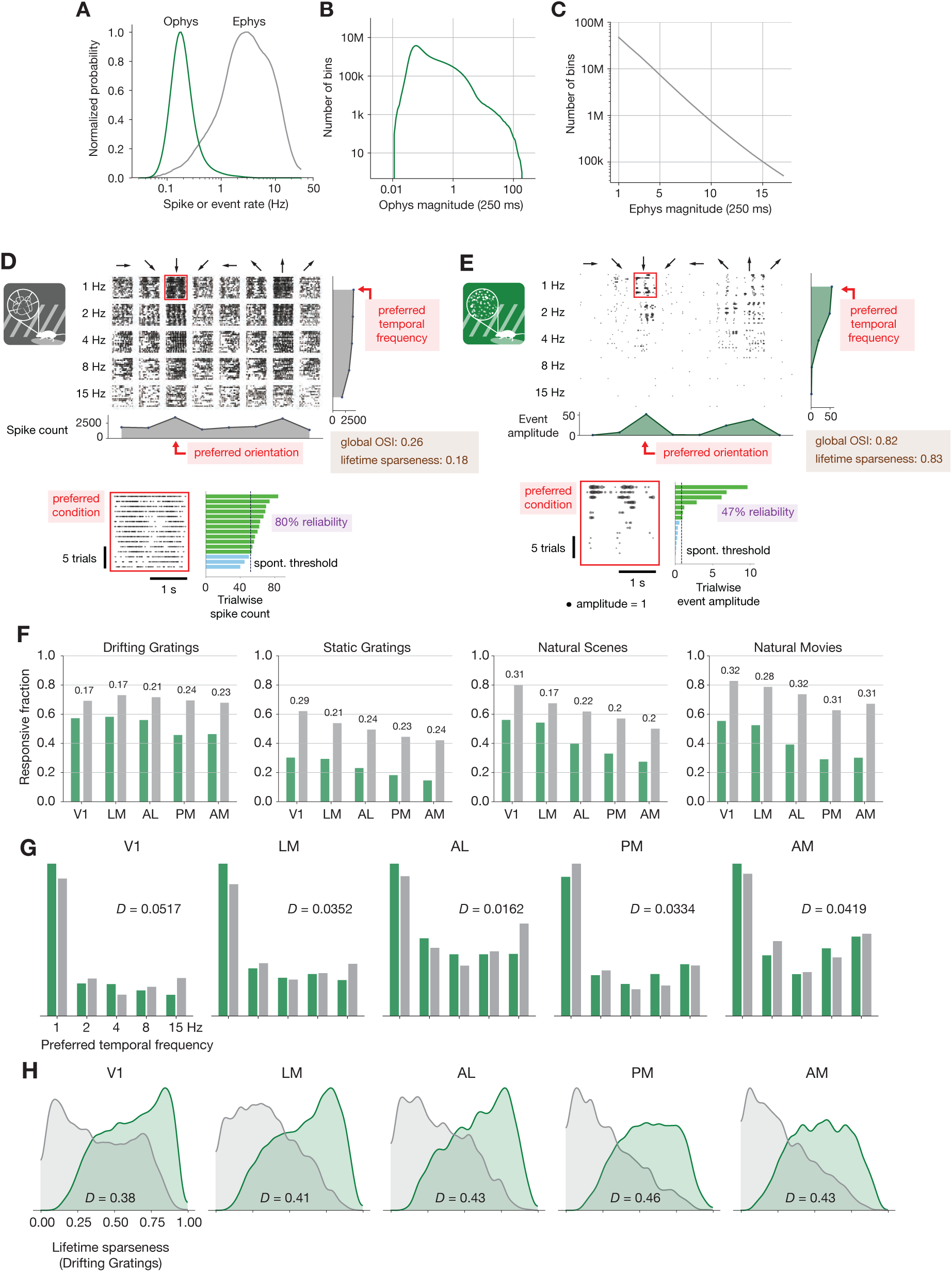
Baseline metric comparison. (A) Normalized histograms of spike or event rates for all ephys (gray) and ophys (green) neurons used in this study. (B) Magnitude of ophys responses for all non-overlapping 250 ms intervals, aggregated over all neurons. Note the log scale on the X- and Y-axes. (C) Magnitudes of ephys responses for all non-overlapping 250 ms intervals, aggregated over all neurons. Note the log scale on the Y-axis only. (D) Drifting gratings spike rasters for an example ephys neuron. Each raster represents 2 s of spikes in response to 15 presentations of a drifting grating at one orientation and temporal frequency. Inset: spike raster for the neuron’s preferred condition, with each trial’s response magnitude shown on the right, and compared to the 95th percentile of the spontaneous distribution. Responsiveness (purple), preference (red), and selectivity (brown) metrics are indicated. (E) Same as D, but for an example ophys neuron. (F) Fraction of neurons deemed responsive to each of four stimulus types, using the same responsiveness metric for both ephys (gray) and ophys (green). Numbers above each pair of bars represent the Jensen-Shannon distance between the full distribution of response reliabilities for each stimulus/area combination. (G) Distribution of preferred temporal frequencies for all neurons in 5 different areas. The value *D* represents the Jensen-Shannon distance between the ephys and ophys distributions. (H) Distributions of lifetime sparseness in response to a drifting grating stimulus for all neurons in 5 different areas. The value *D* represents the Jensen-Shannon distance between the ephys and ophys distributions.

A comparison between individual neurons highlights the effect of these differences on visual physiology. A spike raster from a neuron in V1 recorded with electrophysiology (Figure 2D) appears much denser than the corresponding event raster for a separate neuron that was imaged in the same area (Figure 2E). For each neuron, we computed responsiveness, preference, and selectivity metrics. Both neurons are considered to be responsive to the drifting gratings stimulus class because they have a significant response (*P* < 0.05, compared to a distribution of activity taken during the epoch of spontaneous activity) to the preferred condition (the grating direction and temporal frequency that elicited the largest mean response) on at least 25% of the trials of that condition (response reliability > 25%) (de Vries et al., 2020). Since these neurons were deemed responsive according to this criterion, their function was further characterized in terms of their preferred stimulus condition (the peak of the tuning curve) and their selectivity (a measure of tuning curve sharpness). We use lifetime sparseness (Vinje and Gallant, 2000) as our primary selectivity metric, because it is a general metric that is applicable to every stimulus type. It reflects the distribution of responses of a neuron across some stimulus space (e.g., natural scenes or drifting gratings), equaling 0 if the neuron responds equivalently to all stimulus conditions, and 1 if the neuron only responds to a single condition. Across all areas and mouse lines, lifetime sparseness is highly correlated with more traditional selectivity metrics, such as drifting gratings orientation selectivity (*R* = 0.8 for ephys, 0.79 for ophys; Pearson correlation), static gratings orientation selectivity (*R* = 0.79 for ephys, 0.69 for ophys), and natural scenes image selectivity (*R* = 0.85 for ephys, 0.95 for ophys).

We pooled responsiveness, preference, and selectivity metrics for all of the neurons in a given visual area across experiments, and quantified the disparity between the ophys and ephys distributions using Jensen-Shannon distance. This is the square root of the Jensen-Shannon divergence, which is a method of measuring the similarity between two probability distributions that is symmetric and always has a finite value (Lin, 1991). Jensen-Shannon distance is equal to zero for perfectly overlapping distributions, and one for completely non-overlapping distributions, and falls in between these values for partially overlapping distributions. Across all areas and stimuli, the fraction of responsive neurons was higher in the ephys dataset than the ophys dataset (Figure 2F). To quantify the difference between modalities, we computed the Jensen-Shannon distance for the distributions of response reliabilities, rather than the overall fraction of responsive neurons at the 25% threshold level. While tuning preferences were consistent between the two modalities (Figure 2G), selectivities were much higher in ophys than ephys (Figure 2H). We will continue to refer to these comparisons as “baseline comparisons” in order to distinguish them from subsequent comparisons made after applying one or more transformations to the ophys and/or ephys datasets.

### Effect of laminar sampling bias, running behavior, and transgene expression

As a first step toward reconciling baseline differences in responsiveness and selectivity metrics, we sought to control for high-level variations across the ophys and ephys experimental preparations. For example, the ephys dataset contained more neurons in layer 5, due to the presence of large, highly active somata in this layer. The ophys dataset, on the other hand, had more neurons in layer 4 due to the preponderance of layer 4 Cre lines included in the dataset (Figure S1A). After resampling each dataset to match layer distributions (Figure S1B, see Methods for details), we saw very little change in the overall distributions of responsiveness, preference, and selectivity metrics (Figure S1C-E), indicating that laminar sampling biases are likely not a key cause of the differences we observed.

We next sought to quantify the influence of behavioral differences on our comparison. As running and other motor behavior can influence visually evoked responses (Niell and Stryker, 2010; Stringer et al., 2019a; Vinck et al., 2015; de Vries et al., 2020), could modality-specific behavioral differences contribute to the discrepancies in the response metrics? In our datasets, mice tend to spend a larger fraction of time running in the ephys experiments, perhaps because of the longer experiment duration, which may be further confounded by genotype-specific differences in running behavior (Figure S2A). Within each modality, running had a similar impact on visual response metrics. On average, cells in ephys and ophys have slightly lower responsiveness during periods of running versus non-running (Figure S2B), but slightly higher selectivity (Figure S2C). To control for the effect of running, we sub-sampled ophys experiments in order to match the overall distribution of running fraction to the ephys data (Figure S3A). This transformation had a negligible impact on responsiveness, selectivity, and preference metrics (Figure S3B-D). From this analysis we conclude that, at least for the datasets examined here, behavioral differences do not account for the differences in functional properties inferred from ophys and ephys.

Given that ophys (but not ephys) approaches fundamentally require the expression of exogenous proteins (e.g. Cre, tTA, and GCaMP6 in the case of our transgenic mice) in specific populations of neurons, we sought to determine whether such foreign transgenes, expressed at relatively high levels, could alter the underlying physiology of the neural population. All three proteins have been shown to have neurotoxic effects under certain conditions (Han et al., 2012; Schmidt-Supprian and Rajewsky, 2007; Steinmetz et al., 2017), and calcium indicators, which by design bind intracellular calcium, can additionally interfere with cellular signaling pathways. To examine whether the expression of these genes could explain the differences in functional properties inferred from ophys and ephys experiments, we performed electrophysiology in mice that expressed GCaMP6f under the control of specific Cre drivers. We collected data from mice with GCaMP6f expressed in dense excitatory lines (Cux2 and Slc17a7) or in sparse inhibitory lines (Vip and Sst), and compared the results to those obtained from wild-type mice (Figure 3A). On average, we recorded 45.9 ± 7.5 neurons per area in 17 wild-type mice, and 55.8 ± 15.6 neurons per area in 19 GCaMP6f transgenic mice (Figure 3B). The distribution of firing rates of recorded neurons in mice from all Cre lines was similar to the distribution for units in wild-type mice (Figure 3C).

**Figure 3:**
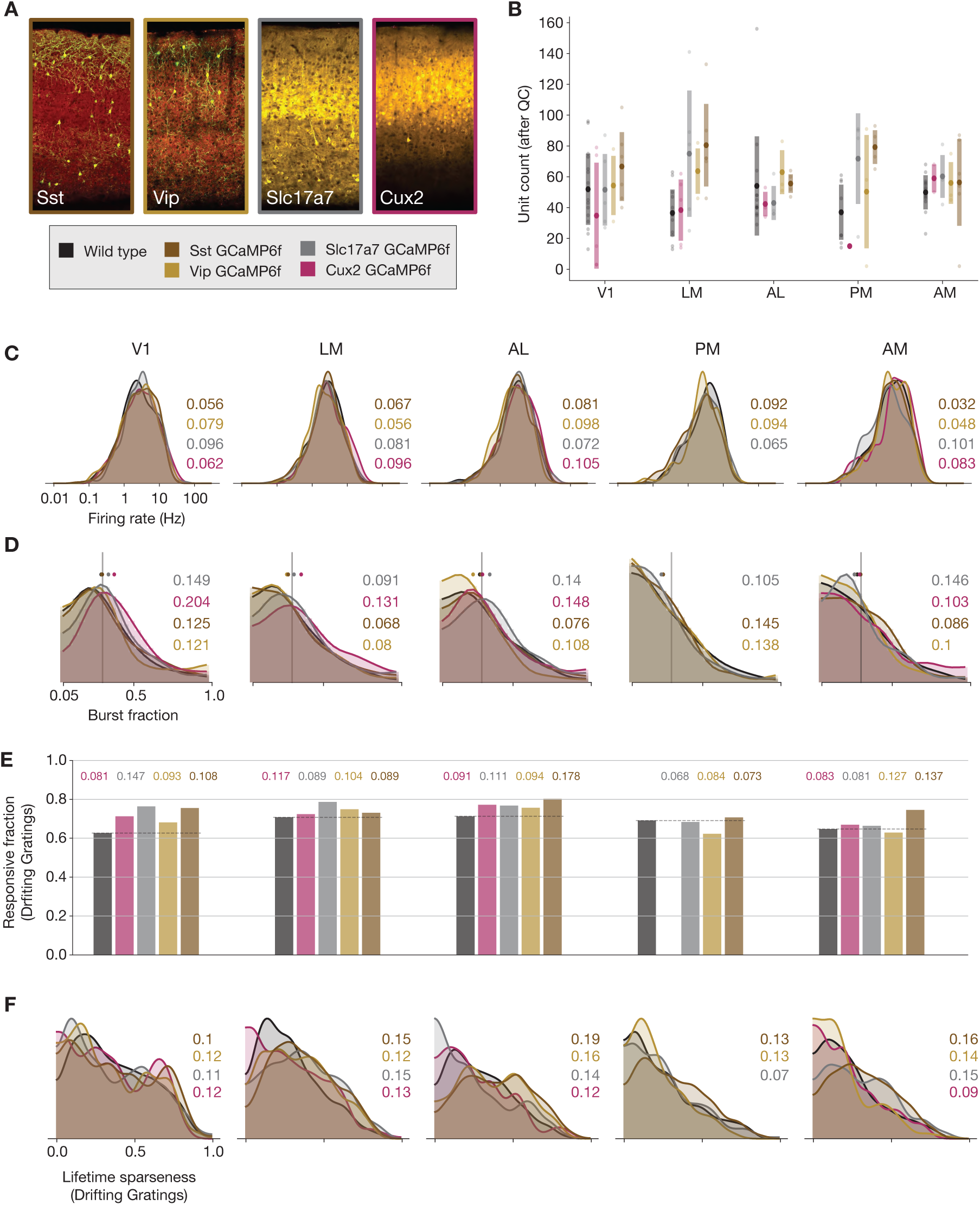
Comparing responses across GCaMP-expressing mouse lines. (A) GCaMP expression patterns for the 4 lines used for ephys experiments, based on two-photon serial tomography (TissueCyte) sections from mice used in the ophys pipeline. (B) Unit yield (following QC filtering) for 5 areas and 5 genotypes. Error bars represent standard deviation across experiments; each dot represents a data point from one experiment. (C) Distribution of firing rates for neurons from each mouse line, aggregated across experiments. (D) Distribution of burst fraction (fraction of all spikes that participate in bursts) for neurons from each mouse line, aggregated across experiments. Dots represent the median of each distribution, shown in relation to a reference value of 0.3. (E) Fraction of neurons deemed responsive to drifting gratings, grouped by genotype. (F) Distribution of lifetime sparseness in response to a drifting grating stimulus, grouped by genotype. In panels (C-F), colored numbers indicate the Jensen-Shannon distance between the wild-type distribution and the distributions of the four GCaMP-expressing mouse lines. Area PM does not include data from Cux2 mice, as it was only successfully targeted in one session for this mouse line.

Because some GCaMP mouse lines have been known to exhibit aberrant seizure-like activity (Steinmetz et al., 2017), we wanted to check whether spike bursts were more prevalent in these mice. We detected bursting activity using the *LogISI* method, which identifies bursts in a spike train based on an adaptive inter-spike interval threshold (Pasquale et al., 2010). The dense excitatory Cre lines showed a slight increase in burst fraction (the fraction of all spikes that participate in bursts) compared to wild-type mice (Figure 3D). This minor increase in burstiness, however, was not associated with changes in responsiveness or selectivity metrics that could account for the baseline differences between the ephys and ophys datasets. The fraction of responsive neurons was not lower in the GCaMP6f mice, as it was for the ophys dataset—in fact, in some visual areas there was an increase in responsiveness in the GCaMP6f mice compared to wild-type (Figure 3E). In addition, the distribution of selectivities was largely unchanged between wild-type and GCaMP6f mice (Figure 3F). Thus, while there may be subtle differences in the underlying physiology of GCaMP6f mice, particularly in the dense excitatory lines, those differences cannot explain the large discrepancies in visual response metrics derived from ephys or ophys.

### Forward-modeling synthetic ophys data from experimental ephys data

Given the substantial differences between the properties of extracellularly recorded spikes and events extracted from fluorescence traces (Figure 2A-C), we hypothesized that transforming spike trains into simulated ophys events could reconcile some of the baseline differences in response metrics we have observed. The inverse transformation—converting ophys events into synthetic spike times—is highly under-specified, due to the reduced temporal resolution (Figure 1B) and event rate (Figure 2A) of ophys. To implement the spikes-to-calcium transformation, we used MLSpike, a biophysically inspired forward model (Deneux et al., 2016). MLSpike explicitly considers the cooperative binding between GCaMP and calcium to generate synthetic ΔF/F fluorescence traces using the spike trains for each cell recorded with ephys as input. We extracted events from these traces using the same *l*_0_-regularized detection algorithm applied to our experimental ophys data, and used these events as inputs to our functional metrics calculations (Figure 4A).

**Figure 4:**
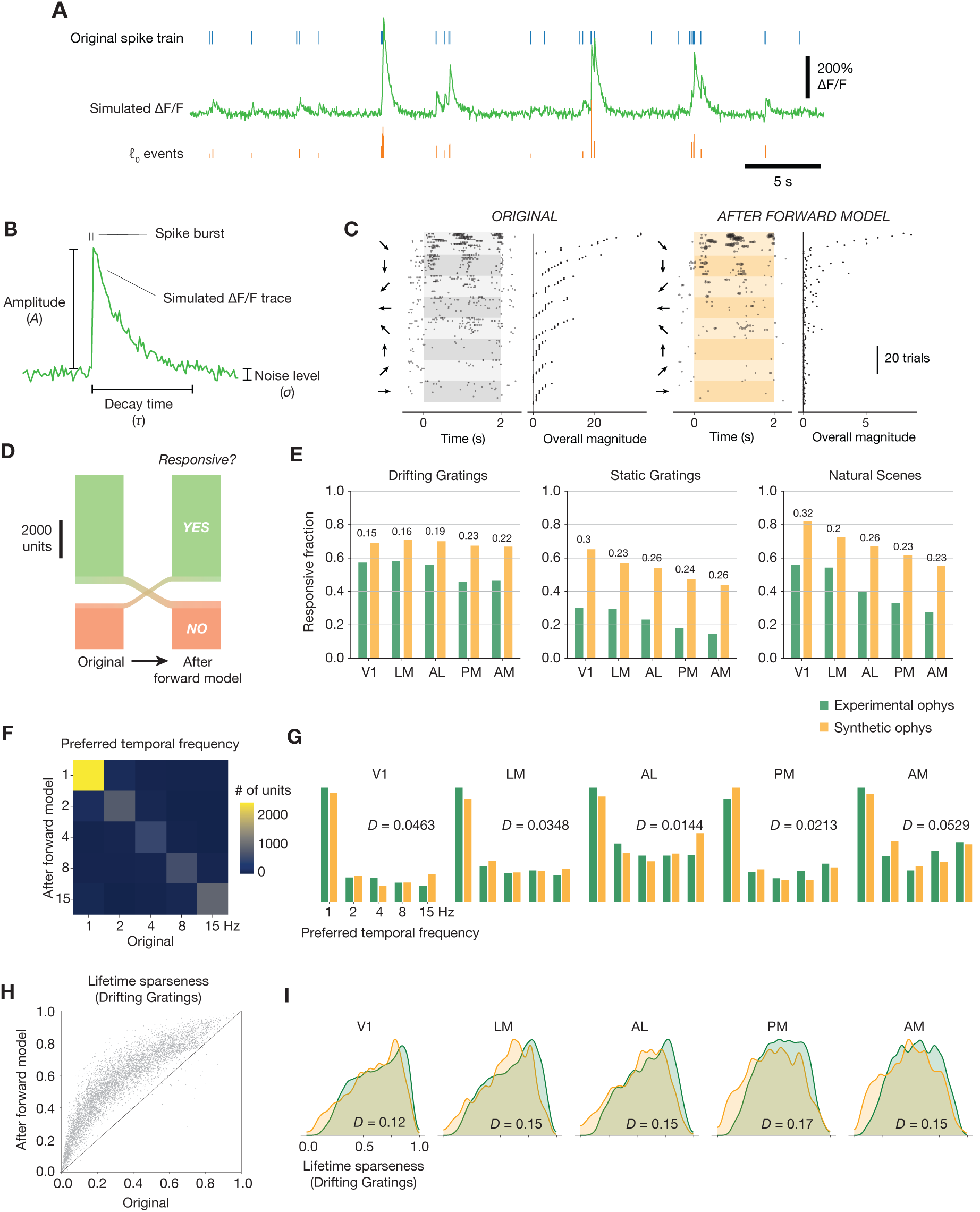
Effects of the spikes-to-calcium forward model. (A) Example of the forward model transformation. A series of spike times (top) is converted to a simulated ΔF/F trace (middle), from which 𝓁_0_-regularized events are extracted using the same method as for the experimental ophys data (bottom). (B) Overview of the three forward model parameters that were fit to data from the Allen Brain Observatory two-photon imaging experiments. (C) Raster plot for one example neuron, before and after applying the forward model. Trials are grouped by drifting grating direction and ordered by response magnitude from the original ephys data. The magnitudes of each trial (based on ephys spike counts or summed ophys event amplitudes) are shown to the right of each plot. (D) Count of neurons responding (green) or not responding (red) to drifting gratings before or after applying the forward model. (E) Fraction of neurons deemed responsive to each of three stimulus types, for both synthetic fluorescence traces (yellow) and true ophys data (green). Numbers above each pair of bars represent the Jensen-Shannon distance between the full distribution of responsive trial fractions for each stimulus/area combination. (F) 2D histogram of neurons’ preferred temporal frequency before and after applying the forward model. (G) Distribution of preferred temporal frequencies for all neurons in 5 different areas. The Jensen-Shannon distance between the synthetic ophys and true ophys distributions is shown for each plot. (H) Comparison between measured lifetime sparseness in response to drifting grating stimulus before and after applying the forward model. (I) Distributions of lifetime sparseness in response to a drifting grating stimulus for all neurons in 5 different areas. The Jensen-Shannon distance between the synthetic ophys and true ophys distributions is shown for each plot.

A subset of the free parameters in the MLSpike model (e.g. ΔF/F rise time, Hill parameter, saturation parameter, and normalized resting calcium concentration) were fit to simultaneously acquired loose patch and two-photon-imaging recordings from layer 2/3 of mouse visual cortex (Huang et al., 2019; Ledochowitsch et al., 2019). Additionally, three parameters were calibrated on the fluorescence traces from the ophys dataset to capture the neuron-to-neuron variance of these parameters: the average amplitude of a fluorescence transient in response to a spike burst (*A*), the decay time of the fluorescence transients (*τ*), and the level of Gaussian noise in the signal (*σ*) (Figure 4B). For our initial characterization, we selected parameter values based on the mode of the overall distribution from the ophys dataset.

The primary consequence of the forward model was to sparsify each neuron’s response by washing out single spikes while non-linearly boosting the amplitude of “burstyßpike sequences with short inter-spike intervals. When responses were calculated on the ephys spike train, a trial containing a 4-spike burst within a 250 ms window would have the same magnitude as a trial with 4 isolated spikes across the 2 s trial. After the forward model, however, the burst would be transformed into an event with a magnitude many times greater than the events associated with isolated spikes, due to the nonlinear relationship between spike counts and the resulting calcium-dependent fluorescence. This effect can be seen in stimulus-locked raster plots for the same neuron before and after applying the forward model (Figure 4C).

What effects does this transformation have on neurons’ inferred functional properties? Applying the forward model to the ephys data did not systematically alter the fraction of responsive cells in the dataset. While 8% of neurons switched from being responsive to drifting gratings to unresponsive, or vice versa, they did so in approximately equal numbers (Figure 4D). The forward model did not improve the match between the distributions of response reliabilities for any stimulus type (Figure 4E). The forward model similarly had a negligible impact on preference metrics; for example, only 14% of neurons changed their preferred temporal frequency after applying the forward model (Figure 4F), and the overall distribution of preferred temporal frequencies still matched that from the ophys experiments (Figure 4G). In contrast, nearly all neurons increased their selectivity after applying the forward model (Figure 4H). Overall, the distribution of lifetime sparseness to drifting gratings became more similar—but still did not completely match—the ophys distribution across all areas (Figure 4I). The average Jensen-Shannon distance between the ephys and ophys distributions was 0.41 before applying the forward model, compared to 0.14 afterward (mean bootstrapped distance between the sub-samples of the ophys distribution = 0.064; *P* = 0.0 for all areas, since bootstrap samples never exceeded the true Jensen-Shannon distance; see Methods for details). These results imply that the primary effects of the forward model—providing a supralinear boost to the “amplitude” of spike bursts, and thresholding out single spike events—can account for baseline differences in selectivity, but not responsiveness, between ephys and ophys.

To assess whether the discrepancies between the ophys and ephys distributions of responsiveness and selectivity metrics could be further reduced by using a different set of forward model parameters, we brute-force sampled 1000 different parameter combinations for one ephys session, using 10 values each for amplitude, decay time, and noise level (Figure S4A), spanning the entire range of parameters calibrated on the experimental ophys data. The fraction of responsive neurons did not change as a function of forward model parameters, except for the lowest values of amplitude and noise level, where it decreased substantially (Figure S4B). This parameter combination (*A ≤* 0.0015, *σ ≤* 0.03) was observed in less than 1% of actual neurons recorded with ophys, so it cannot account for differences in responsiveness between the two modalities. Both the difference between the median lifetime sparseness for ophys and ephys, as well as the Jensen-Shannon distance between the full ephys and ophys lifetime sparseness distributions, were near the global minimum for the parameter values we initially used (Figure S4C,D).

It is conceivable that the inability of the forward model to fully reconcile differences in responsiveness and selectivity was due to the fact that we applied the same parameters across all neurons of the ephys dataset, without considering their genetically defined cell type. To test for cell-type specific differences in forward model parameters, we examined the distributions of amplitude, decay time, and noise level for individual excitatory Cre lines used in the ophys dataset. The distributions of parameter values across genotypes were largely overlapping, with the exception of increasing noise levels for some of the deeper populations (e.g. Tlx3-Cre in layer 5, and Ntsr1-Cre GN220 in layer 6) and an abundance of low-amplitude cells in the Fezf2-CreER population (Figure S5A). Given that higher noise levels and lower amplitudes did not improve the correspondence between the ephys and ophys metric distributions, we concluded that selecting parameter values for individual neurons based on their most likely cell type would not change our results. Furthermore, we saw no correlation between responsiveness or selectivity metrics in ophys neurons and their calibrated amplitude, decay time, or noise level (Figure S5B-E). This implies that there is no underlying relationship between forward model parameters and functional properties in the experimental data.

### Effect of ephys selection bias

We next sought to determine whether electrophysiology’s well-known selection bias in favor of more active neurons could account for the differences between modalities. Whereas calcium imaging can detect the presence of all neurons in the field of view that express a fluorescent indicator, ephys cannot detect neurons unless they fire action potentials. This bias is exacerbated by the spike sorting process, which requires a sufficient number of spikes in order to generate an accurate template of each neuron’s waveform. Spike sorting algorithms can also mistakenly merge spikes from nearby neurons into a single “unit” or allow background activity to contaminate a spike train, especially when spike waveforms generated by one neuron vary over time, e.g. due to the adaptation that occurs during a burst. These issues all result in an apparent activity level increase in ephys recordings. Indeed, intracellular recordings in L2/3 of awake mouse V1 report average spontaneous firing rates well below 1 Hz (Einstein et al., 2017; Haider et al., 2013), substantially lower than the average rate of 2.2 Hz measured in L2/3 of our dataset. In addition, assuming a 50-micron “listening radius” for the probes (radius of half-cylinder around the probe where the neurons’ spike amplitude is sufficiently above noise to trigger detection) (Buzsáki, 2004; Harris et al., 2016; Shoham et al., 2006), the average yield of 116 regular-spiking units/probe (prior to QC filtering) would imply a density of 42,000 neurons/mm^3^, much lower than the known density of *∼*90,000 neurons/mm^3^ for excitatory cells in mouse visual cortex (Erö et al., 2018).

If the ephys dataset is biased toward recording neurons with higher firing rates, it may be more appropriate to compare it with only the most active neurons in the ophys dataset. To test this, we systematically increased the event rate threshold for the ophys neurons, so the remaining neurons used for comparison were always in the upper quantile of mean event rate. Applying this filter increased the overall fraction of responsive neurons in the ophys dataset, such that the experimental ophys and synthetic ophys distributions had the highest similarity when between 7-39% of the most active ophys neurons were included (V1: 39%, LM: 34%, AL: 25%, PM: 7%, AM: 14%) (Figure 5A). This indicates that more active neurons tend to be more responsive to our visual stimuli, which could conceivably account for the discrepancy in overall responsiveness between the two modalities. However, applying this event rate threshold actually increased the differences between the selectivity distributions, as the most active ophys neurons were also more selective (Figure 5B). Thus, sub-selection of ophys cells based on event rate was not sufficient to fully reconcile the differences between ephys and ophys.

**Figure 5:**
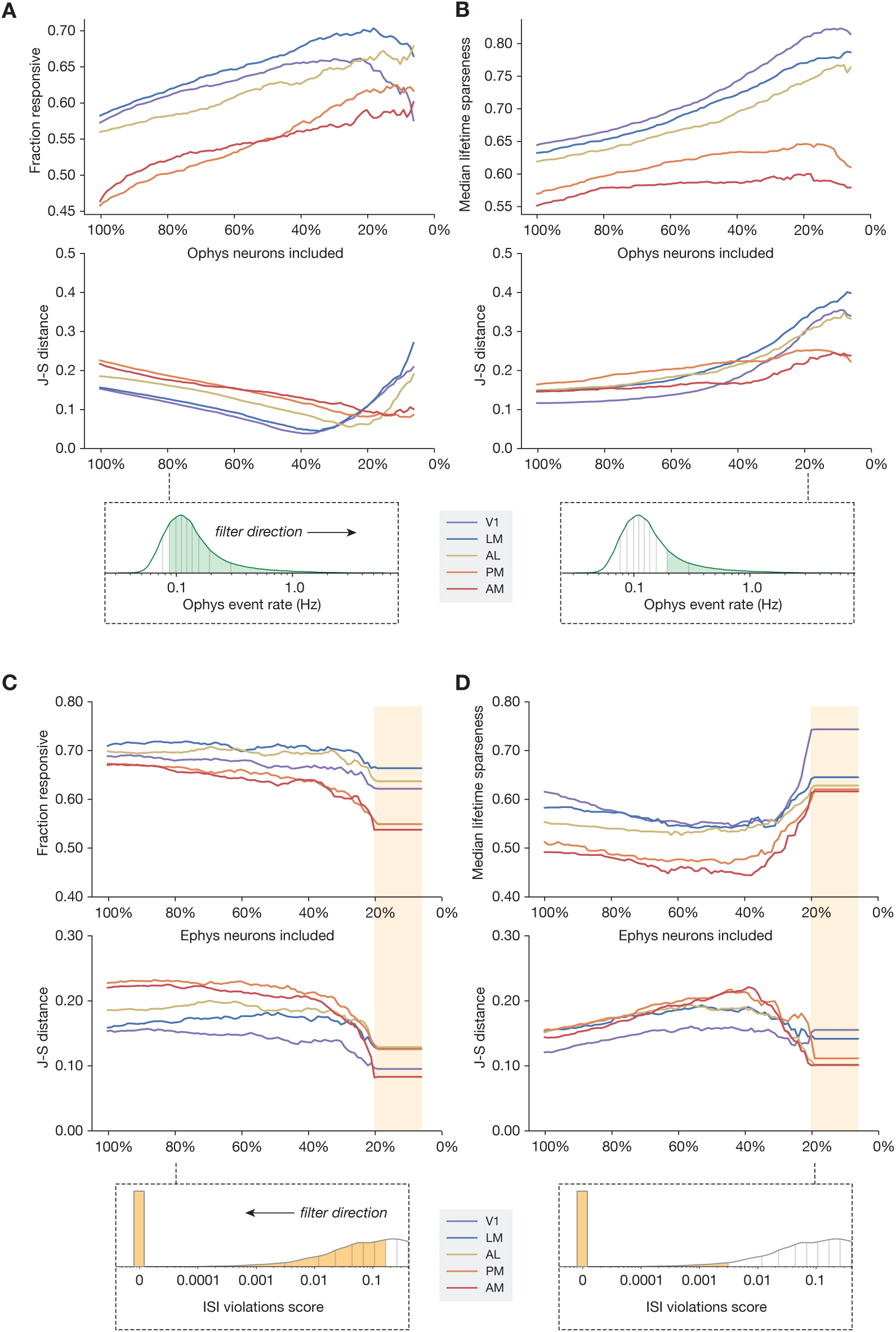
Sub-sampling based on event rate or ISI violations. (A) *Top:* Change in fraction of neurons responding to drifting gratings for each area in the ophys dataset, as function of the percent of ophys neurons included in the comparison. *Middle:* Jensen-Shannon distance between the synthetic ophys and true ophys response reliability distributions. *Bottom:* Overall event rate distribution in the ophys dataset. As the percent of included neurons decreases (from left to right), more and more neurons with low activity rates are excluded. (B) Same as A, but for drifting gratings lifetime sparseness (selectivity) metrics. (C) Top: Change in fraction of neurons responding to the drifting gratings stimulus for each area in the ephys dataset, as function of the percent of ephys neurons included in the comparison. Middle: Jensen-Shannon distance between the synthetic ophys and true ophys response reliability distributions. Bottom: Overall ISI violations score distribution in the ephys dataset. As the percent of included neurons decreases (from right to left), more and more neurons with high contamination levels are excluded. Yellow shaded area indicates the region where the minimum measurable contamination level (ISI violations score = 0) is reached. (D) Same as C, but for drifting gratings lifetime sparseness (selectivity) metrics.

If the ephys dataset includes spike trains that are contaminated with spurious spikes from one or more nearby neurons, then it may help to compare our ophys results only to the least contaminated neurons from the ephys dataset. Our initial QC process excluded cells with an inter-spike interval (ISI) violations score (Hill et al., 2011) above 0.5, to remove highly contaminated units, but while the presence of refractory period violations implies contamination, the absence of such violations does not imply error-free clustering, so some contamination may remain. We systematically decreased our tolerance for ISI-violating ephys neurons, so the remaining neurons used for comparison were always in the lower quantile of contamination level. For the most restrictive thresholds, where there was zero detectable contamination in the original spike trains (ISI violations score = 0), the match between the synthetic ophys and experimental ophys selectivity and responsiveness distributions was maximized (Figure 5C,D). This indicates that, unsurprisingly, contamination by neighboring neurons (as measured by ISI violations score) reduces selectivity and increases responsiveness. Therefore, the inferred functional properties are most congruent across modalities when the ephys analysis includes a stringent threshold on the maximum allowable contamination level.

## Results across stimulus types

The previous results have primarily focused on the drifting gratings stimulus, but we observe similar effects for all of the stimulus types shared between the ophys and ephys datasets. Figure 6 summarizes the impact of each transformation we performed, either before or after applying the forward model, for drifting gratings, static gratings, natural scenes, and natural movies.

**Figure 6:**
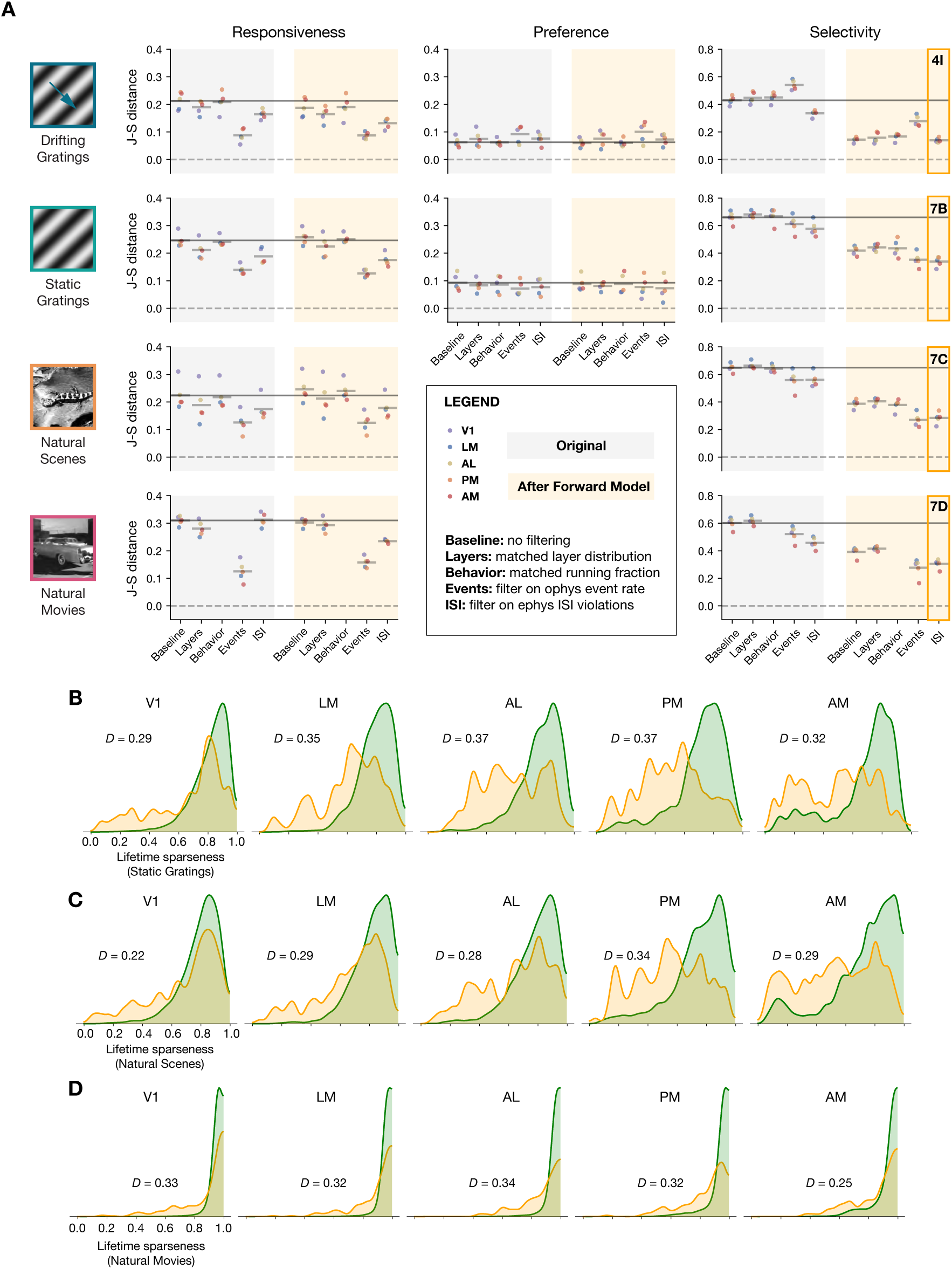
Summary of results for all stimulus types. (A) Jensen-Shannon distance between the ephys and ophys distributions for three metrics and four stimulus classes. Individual dots represent visual areas, and horizontal lines represent the mean across areas. Each column represents a different post-processing step, with comparison performed on either the original ephys data (gray), or the synthetic fluorescence data obtained by passing experimental ephys data through a spikes-to-calcium forward model (yellow). (B) Full distributions of lifetime sparseness in response to static gratings for experimental ophys (green) and synthetic ophys (yellow). (C) same as B, but for natural scenes. (D), same as B and C, but for natural movies. The distribution for drifting gratings is shown in Figure 4I.

Across all stimulus types, the forward model had very little impact on responsiveness. Instead, subselecting the most active neurons from ophys using an event-rate filter rendered the shape of the distributions the most similar. For the stimulus types for which we could measure preference across small number of categories (temporal frequency of drifting gratings and spatial frequency of static gratings), no data transformations were able to improve the overall match between the ephys and ophys distributions, as they were already very similar in the baseline comparison. For selectivity metrics (lifetime sparseness), applying the forward model played the biggest role in improving cross-modal similarity, although there was a greater discrepancy between the resulting distributions for static gratings, natural scenes, and natural movies than there was for drifting gratings. Filtering ephys neurons based on ISI violations score further reduced the Jensen-Shannon distance, but it still remained well above zero. This indicates that the transformations we employed could not fully reconcile observed differences in selectivity between ephys and ophys.

### Classifying neurons based on response reliability

The differences in responsiveness and selectivity metrics computed from the ephys and ophys datasets highlight that functional properties of neurons in the mouse visual cortex can appear to be dependent on the choice of recording modality. How might these differences affect attempts to group neurons into functional clusters? Previously, a functional classification scheme based on responses to all four stimulus types revealed distinct response classes in the ophys dataset (de Vries et al., 2020). The largest class contained neurons that did not respond reliably to any of the stimuli, while only *∼*10% of neurons responded reliably to all stimuli. This classification reveals that many neurons in the visual cortex are not well described by classical models, and may represent or integrate more complex visual and non-visual features. We sought to test whether this classification would be impacted by modality-specific biases.

Using a Gaussian mixture model, we performed unsupervised clustering on the 4 x *N* matrix of response reliabilities for each neuron’s preferred condition for each stimulus class (drifting gratings, static gratings, natural scenes, and natural movies), where *N* represents the number of neurons in the ephys or ophys datasets (Figure 7A). The resulting clusters were given class labels based on whether their mean response reliability was above an initial threshold of 25% (to match what we used in our previous analysis of responsiveness). After labeling each cluster, we calculated the fraction of neurons belonging to each functional class. We repeated this clustering process 100 times with different initialization conditions, to yield the average class membership of neurons across runs (Figure 7B).

**Figure 7:**
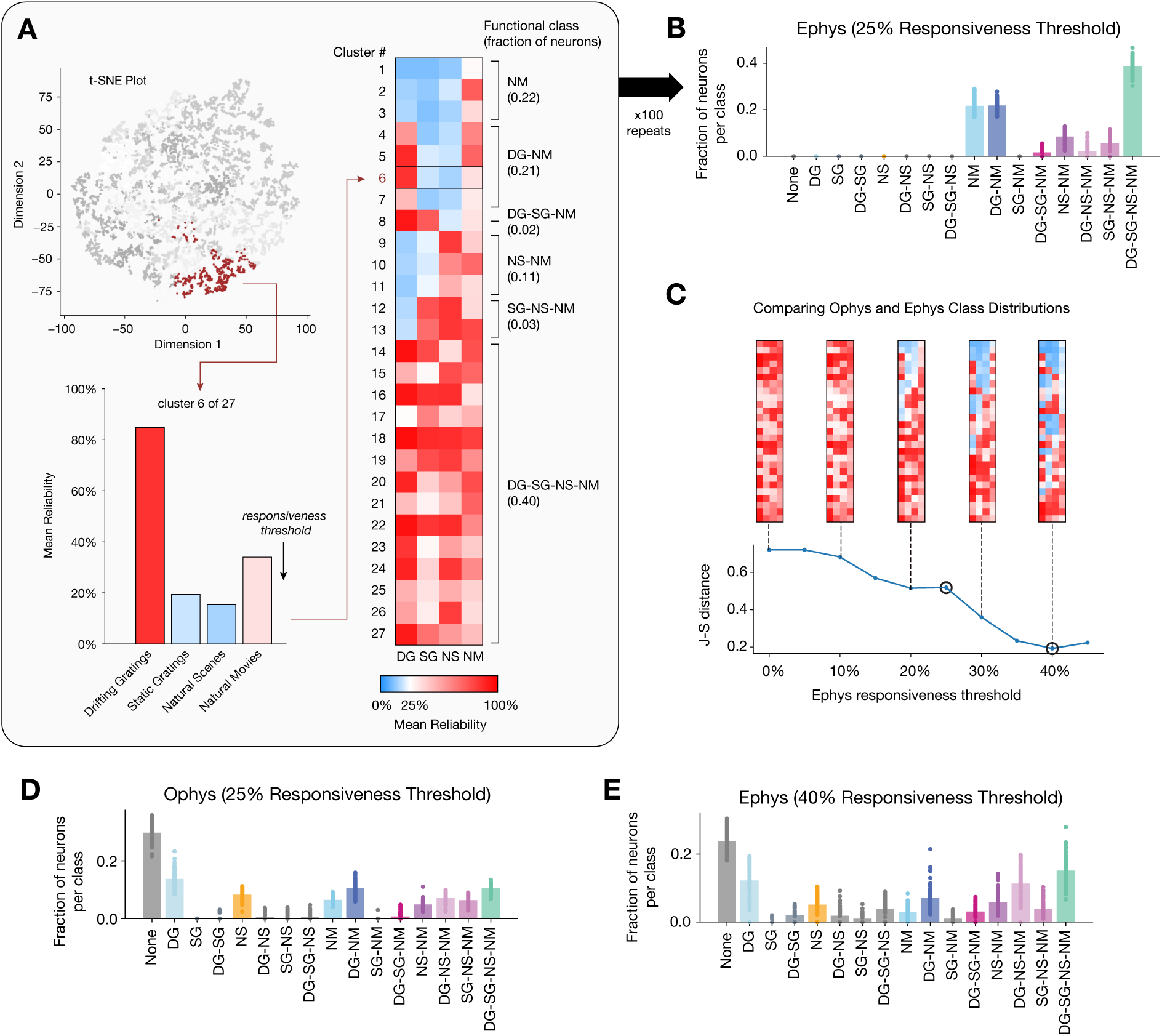
Identifying functional classes based on response reliability. (A) Example clustering run for all neurons in the ephys dataset. First, unsupervised clustering is performed on the matrix of response reliabilities to four stimulus types. Colors on the t-SNE plot represent different cluster labels, with one cluster highlighted in brown. Next, the neurons in each cluster are used to compute the mean reliability for each stimulus type. The mean reliabilities for all 27 clusters are shown in the heatmap, with the 25% responsiveness threshold indicated by the transition from blue to red. Each cluster is assigned to a class based on the stimulus types for which its mean reliability is above threshold. (B) Strip plots representing the fraction of neurons belonging to each class, averaged over 100 repeats of the Gaussian mixture model for ephys (*N* = 11,030 neurons), at a 25% threshold level. (C) Jensen-Shannon distance between the ephys and ophys class membership distributions, as a function of the ephys responsiveness threshold. The points plotted in panels B and E are circled. Heat maps show the mean reliabilities relative to threshold for all 27 clusters shown in panel A. (D) Strip plots representing the fraction of neurons belonging to each class, averaged over 100 repeats of the Gaussian mixture model for ophys (*N* = 20,611 neurons), at a 25% threshold level. (E) Same as B, but for the 40% threshold level that minimizes the difference between the ophys and ephys distributions.

For ephys, the largest class (with *∼*40% of the neurons) contains neurons that were above threshold for all four classes of stimuli, i.e. the neurons responded reliably to all stimulus types. Two other classes contained the bulk of the remaining neurons, in which the neurons responded to natural movies, or to both drifting gratings and natural movies. For ophys, on the other hand, the largest class was that in which the neurons were above threshold for none of the stimuli, consistent with previous work (de Vries et al., 2020) (Figure 7D). A handful of other classes had *∼*10% of the neurons, including the classes in which neurons responded to drifting gratings, both drifting gratings and natural movies, and all four stimuli. Thus, naively applying the same analysis to both modalities revealed very different landscapes of functional classes.

The fact that many more neurons appear to respond to all stimulus types in the ephys dataset is consistent with the fact that responsiveness is higher in the ephys dataset for each stimulus (Figure 2F), likely reflecting the bias of electrophysiology towards recording more active units (Figure 5A). To account for this, we systematically raised the threshold used to group clusters into functional classes for the ephys dataset and found that the distribution of classes became more similar to the distribution of classes for the ophys dataset (Figure 7C). A threshold of 40% response reliability for the ephys dataset minimizes the Jensen-Shannon distance between the distributions, making the class assignments in ephys look remarkably similar to those for the ophys dataset (Figure 7E). We consider the class labels for each neuron an indication of its “meta-preference” for different stimulus types. Thus, once we account for the higher levels of responsiveness seen in the ephys dataset, we observe similar meta-preferences to the ophys dataset. This is consistent with the prediction—based on our finding that the forward model largely conserves neuronal tuning—that the overall preference for particular stimulus conditions (e.g. temporal frequencies) should be largely invariant across the two recording modalities (Figure 2G).

## Discussion

In this study, we have compared response metrics derived from mouse visual cortex excitatory populations collected under highly standardized conditions, but using two different recording modalities. Overall, we observe similar stimulus preferences across the two datasets (e.g. preferred temporal frequencies within each visual area), but we see systematic differences in responsiveness and selectivity. Prior to any attempt at reconciliation, electrophysiological recordings showed a higher fraction of cells with stimulus-driven activity, while calcium imaging showed higher selectivity (sharper tuning) among responsive neurons.

Leveraging our understanding of the spikes-to-calcium forward model successfully accounted for some, but not all, of these discrepancies. Notably, the forward model boosted selectivity due to the sparsifying effect of the calcium dynamics. As large, burst-dependent calcium transients can have amplitudes several orders of magnitude above the median amplitude (Figure 2B), this causes the response to the preferred stimulus condition to be weighted more heavily in selectivity calculations than non-preferred conditions. Similarly, isolated spikes during non-preferred conditions can be virtually indistinguishable from noise when viewed through the lens of the forward model. When the same trials are viewed through the lens of electrophysiology, however, spike counts increase more or less linearly, leading to the appearance of lower selectivity.

Unexpectedly, the forward model did not change responsiveness metrics. We initially hypothesized that the lower responsiveness in ophys was due to the fact that single-spike events are often not translated into detectable calcium transients (Ledochowitsch et al., 2019). Instead, our observation of unchanged responsiveness following the forward model suggests that differences between modalities are more likely due to ephys sampling bias—i.e., extracellular recordings missing small or low-firing rate cells, or merging spike trains from nearby cells. In order to reconcile some of these differences, we could either apply a very strict threshold on ISI violations score to the ephys dataset, or remove between 61-93% of the least active neurons from the ophys dataset. This should serve as a cautionary tale for anyone estimating the fraction of neurons in an area that appear to increase their firing rate in response to environmental or behavioral events. Without careful controls, contamination from other cells in the vicinity can make this fraction appear artificially high. While it may be possible to minimize these impurities with improved spike-sorting algorithms, there will always be neurons that even the best algorithms will not be able to distinguish in the face of background noise.

Differences in laminar sampling and running behavior between the two modalities had almost no effect in our comparisons. The transgenic expression of GCaMP6f in specific neural populations also did not impact the distributions of functional metrics. Finally, the initial parameters chosen for the forward model produced metric distributions that were close to the optimum match over the realistic parameter space. Therefore, we conclude that the primary contribution to differences in the considered ephys and ophys metrics comes from (1) the intrinsic nature of the spikes-to-calcium transfer function (2) the selection bias of extracellular electrophysiology recordings.

Even after accounting for these known factors to the best of our ability, the overall ophys population still displayed higher selectivity than its ephys counterpart (Figure 4I, Figure 6B-D). What could account for these remaining differences? One possibility is that there may be residual undetected contamination in our ephys recordings. An ISI violations score of 0 does not guarantee that there is no contamination, just that we are not able to measure it using this metric. Sampling the tissue more densely (i.e. increasing the spatial resolution of spike waveforms) or improving spike sorting methods could reduce this issue. Another possibility is that “missed” spikes—especially those at the end of a burst—could result in reduced amplitudes for the simulated calcium transients. In addition, if the *in vivo* spikes-to-calcium transfer function is non-stationary, there could be stimulus-dependent changes in calcium concentration that are not captured by a forward model that takes a spike train as its only input. Simultaneous cell-attached electrophysiology and two-photon imaging experiments have demonstrated the existence of “prolonged depolarization events” (PDEs) in some neurons that result in very large increases in calcium concentration, but that are indistinguishable from similar burst events in extracellular recordings (Ledochowitsch et al., 2019).

The forward models available in the literature (Deneux et al., 2016; Wei et al., 2019) are of comparable power, in that their most complex instantiations allow for non-instantaneous fluorescence rise as well as for a non-linear relationship between calcium concentration and fluorescence. To the best of our knowledge, none of these forward models explicitly model non-stationarities in the spike-to-calcium transfer function. Moreover, all currently available models suffer from the drawback that fits to simultaneously recorded ground truth data yield significant variance in model parameters across neurons (Ledochowitsch et al., 2019; Wei et al., 2019). We strove to mitigate this shortcoming by showing that brute-force exploration of MLSpike model parameter space could not significantly improve the match between real and synthetic ophys data Figure S4.

While results obtained from ephys and ophys are sometimes considered interchangeable from a scientific standpoint, in actuality they each provide related but fundamentally different perspectives on the underlying neural activity. Extracellular electrophysiology tells us—with sub-millisecond temporal resolution— about a neuron’s spiking outputs. Calcium imaging doesn’t measure the outgoing action potentials, but rather their impact on a neuron’s internal state, in terms of increases in calcium concentration that drive various downstream pathways (West et al., 2001). While voltage-dependent fluorescent indicators may offer the best of both worlds, there are substantial technical hurdles to employing them on comparably large scales (Peterka et al., 2011). Thus, in order to correctly interpret existing and forthcoming datasets, we must account for the inherent biases of these two recording modalities.

A recent study comparing matched populations recorded with electrophysiology or imaging emphasized differences in the temporal profiles of spike trains and calcium-dependent fluorescence responses (Wei et al., 2019). The authors found that event-extraction algorithms that convert continuous ΔF/F traces to putative spike times could not recapitulate the temporal profiles measured with electrophysiology; on the other hand, a novel forward model that transformed spike times to synthetic ΔF/F traces could make their electrophysiology results appear more like those from the imaging experiments. Their conclusions were primarily based on metrics derived from the evolution of firing rates or ΔF/F ratios over the course of a behavioral trial. However, there are other types of functional metrics that are not explicitly dependent on temporal factors, such as responsiveness and selectivity, which cannot be reconciled using a forward model alone.

More work is needed to understand the detailed physiological underpinning of the modality-specific differences we have observed. One approach, currently underway at the Allen Institute and elsewhere, is to carry out recordings with extremely high-density silicon probes (Neuropixels Ultra), up to 20x higher than the Neuropixels probes used in this study. Such probes can capture each spike waveform with 100 or more electrodes, making it easier to disambiguate waveforms from nearby cells, and making it less likely that neurons with small somata would evade detection or that their waveforms would be mistaken for those of other units. These experiments should make it easier to quantify the selection bias of extracellular electrophysiology, as well as the degree to which missed cells and contaminated spike trains have influenced the results of the current study. In addition, experiments in which silicon probes are combined with two-photon imaging—either through interleaved sampling, spike-triggered image acquisition, or improved artifact removal techniques—could provide ground-truth information about the relationship between extracellular electrophysiology and calcium imaging. Due to the technical difficulties of such experiments, it is unlikely that the yield would be very high, so we expect that these will serve to complement and validate our approach of comparing large-scale unimodal datasets.

Ultimately, the goal of this work is not to determine which of these two methods is “best,” since their complementary strengths ensure they will each remain essential to scientific progress for many years to come. Instead, we want to establish guidelines for properly interpreting the massive amounts of data that have been or will be collected using either modality. From this study, we have learned that extracellular electrophysiology overestimates the fraction of neurons that elevate their activity in response to visual stimuli, in a manner that is consistent with the effects of selection bias and contamination. Further, the apparent differences in selectivity underscore the fact that we need to consider what each modality is actually measuring, as selectivity metrics based on spike counts (the neuron’s outputs) will almost always be lower than selectivity metrics based on calcium concentrations (the neuron’s internal state). Even with this in mind, however, we cannot fully reproduce the observed levels of calcium-dependent selectivity using spike times alone—suggesting that a neuron’s internal state may contain stimulus-specific information that is not necessarily reflected in its outputs.

## Acknowledgements

We thank the Allen Institute founder, Paul G. Allen, for his vision, encouragement, and support. We thank the MindScope Scientific Advisory Council for providing feedback on our initial results. We thank the TissueCyte team, including Nadezhda Dotson, Mike Taormina, and Anh Ho, for histology and imaging associated with Figure 3A. We thank Adam Charles, Jeff Gauthier, and David Tank for helpful discussions on data processing methods. We thank Nicholas Cain for developing the computational infrastructure that made this project feasible, as well as for his enthusiastic analysis of pilot data in the early stages of this project. We thank Corbett Bennett and Jun Zhuang for their feedback on the manuscript.

## Author Contributions

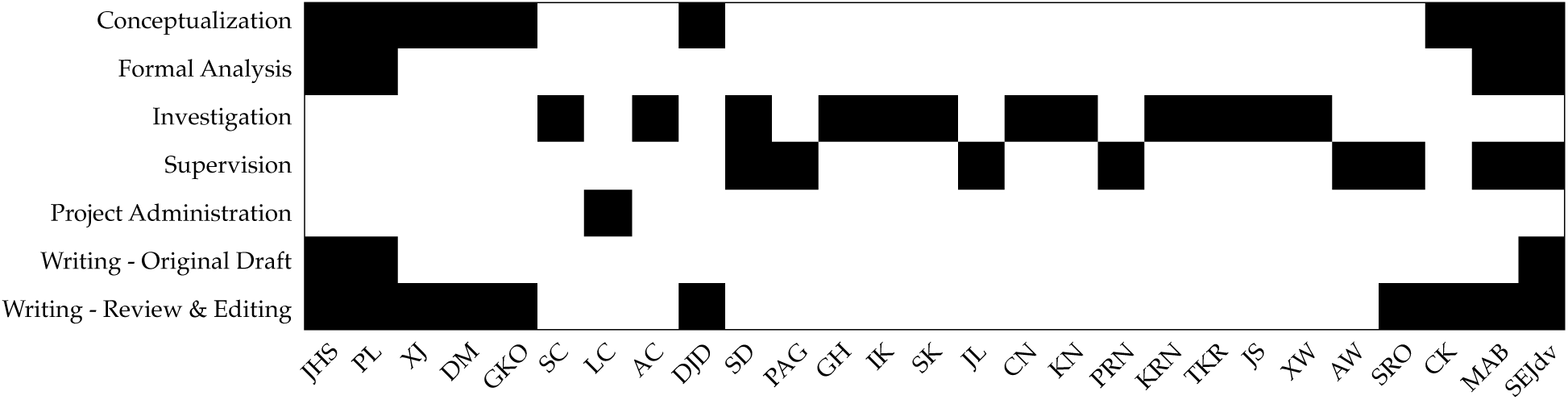

## Methods

### Previously Released Data

We used two-photon calcium imaging recordings from the Allen Brain Observatory Visual Coding dataset (de Vries et al., 2020; ©2016 Allen Institute for Brain Science, available from observatory.brain-map.org). This dataset consists of calcium fluorescence time series from 63,521 neurons in 6 different cortical areas across 14 different transgenic lines. Neurons were imaged for three separate sessions (A, B, and C), each of which used a different visual stimulus set (Figure 1E). Our analysis was limited to neurons in 5 areas (V1, LM, AL, PM, and AM) and 10 lines expressing GCaMP6f in excitatory cells, and which were present in either session A, session B, or both (total of 41,578 neurons).

We used extracellular electrophysiological recordings from the Allen Brain Observatory Neuropixels dataset (Siegle et al., 2019; ©2019 Allen Institute for Brain Science, available from portal.brain-map.org). This dataset consists of spike trains from 99,180 “units” (putative neurons with varying degrees of completeness and contamination) from 58 mice in a variety of cortical and subcortical structures. We limited our analysis to 31 sessions that used the “Brain Observatory 1.1” stimulus set (Figure 1E) and units (hereafter, “neurons”) from 5 visual cortical areas (V1, LM, AL, PM, and AM) that displayed “regular spiking” action potential waveforms (peak-to-trough interval > 0.4 ms). Only neurons that passed the following quality control thresholds were included: presence ratio > 0.9 (fraction of the recording session during which spikes are detected), amplitude cutoff < 0.1 (estimate of the fraction of missed spikes), and ISI violations score < 0.5 (Hill et al., 2011) (estimate of the relative rate of contaminating spikes). After these filtering steps, there were 5917 neurons for analysis.

### Neuropixels Recordings in GCaMP6f Mice

We collected a novel electrophysiology dataset from transgenic mice expressing GCaMP6f, as well as additional wild-type mice. Experimental procedures closely followed those described in Siegle et al., 2019, and are summarized below.

Mice were maintained in the Allen Institute animal facility and used in accordance with protocols approved by the Allen Institute’s Institutional Animal Care and Use Committee. Five genotypes were used: wild-type C57BL/6J mice purchased from Jackson Laboratories (*N* = 2) or Vip-IRES-Cre;Ai148 (*N* = 3), Sst-IRES-Cre;Ai148 (*N* = 6), Slc17a7-IRES2-Cre;Camk2a-tTA;Ai93 (*N* = 5), and Cux2-CreERT2;Camk2a-tTA;Ai93 (*N* = 5) mice bred in-house. Following surgery, mice were single-housed and maintained on a reverse 12-hour light cycle. All experiments were performed during the dark cycle.

At around age P80, mice were implanted with a titanium headframe. In the same procedure, a 5 mm diameter piece of skull was removed over visual cortex, followed by a durotomy. The skull was replaced with a circular glass coverslip coated with a layer of silicone to reduce adhesion to the brain surface.

On the day of recording (at least four weeks after the initial surgery), the glass coverslip was removed and replaced with a plastic insertion window containing holes aligned to six cortical visual areas, identified via intrinsic signal imaging (Garrett et al., 2014). An agarose mixture was injected underneath the window and allowed to solidify. This mixture was optimized to be firm enough to stabilize the brain with minimal probe drift, but pliable enough to allow the probes to pass through without bending. At the end of this procedure, mice were returned to their home cages for 1-2 hours prior to the recording session.

All recordings were carried out in head-fixed mice using Neuropixels 1.0 probes (Jun et al., 2017; available from neuropixels.org) mounted on 3-axis stages from New Scale Technologies (Victor, NY). These probes have 383 recording sites oriented in a checkerboard pattern on a 70 µm wide x 10 mm long shank, with 20 µm vertical spacing. Data streams from each electrode were acquired at 30 kHz (spike band) and 2.5 kHz (LFP band) using the Open Ephys GUI (Siegle et al., 2017). Gain settings of 500x and 250x were used for the spike band and LFP band, respectively. Recordings were referenced to a large, low-impedance electrode at the tip of each probe.

Pre-processing, spike sorting, and quality control methods were identical to those used for the previously released dataset (code available at github.com/alleninstitute/ecephys spike sorting and github.com/mouseland/kilosort2). Filtering by brain region (V1, LM, AL, PM, and AM), waveform width (>0.4 ms), and QC metrics (presence ratio > 0.9, amplitude cutoff < 0.1, ISI violations score < 0.5) yielded 5113 neurons for analysis. For all analyses except for those in Figure 3, neurons from this novel dataset were grouped with those from the previously released dataset, for a total of 11,030 neurons.

Neurons were registered to 3D brain volumes obtained with an open-source optical projection tomography system (github.com/alleninstitute/AIBSOPT). Brains were first cleared using a variant of the iDISCO method (Renier et al., 2014), then imaged with white light (for internal structure) or green light (to visualize probe tracks labeled with fluorescent dye). Reconstructed volumes were mapped to the Mouse Common Coordinate Framework (CCFv3) (Wang et al., 2020) by matching key points in the original brain to corresponding points in a template volume. Finally, probes tracks were manually traced and warped into the CCFv3 space, and electroes were aligned to structural boundaries based on physiological landmarks (Siegle et al., 2019).

### Visual Stimuli

Analysis was limited to epochs of drifting gratings, static gratings, natural scenes, or natural movie stimuli, which were shown with identical parameters across the two-photon imaging and electrophysiology experiments. Visual stimuli were generated using custom scripts based on PsychoPy (Peirce, 2007) and were displayed using an ASUS PA248Q LCD monitor, 1920 × 1200 pixels in size (21.93” wide, 60 Hz refresh rate). Stimuli were presented monocularly, and the monitor was positioned 15 cm from the mouse’s right eye and spanned 120° x 95° of visual space prior to stimulus warping. Each monitor was gamma corrected and had a mean luminance of 50 cd/m^2^. To account for the close viewing angle of the mouse, a spherical warping was applied to all stimuli to ensure that the apparent size, speed, and spatial frequency were constant across the monitor as seen from the mouse’s perspective.

The drifting gratings stimulus consisted of a full-field sinusoidal grating at 80% contrast presented for 2 seconds, followed by a 1 s mean luminance gray period. Five temporal frequencies (1, 2, 4, 8, 15 Hz), eight different directions (separated by 45°), and one spatial frequency (0.04 cycles per degree) were used. Each grating condition was presented 15 times in random order.

The static gratings stimulus consisted of a full field sinusoidal grating at 80% contrast that was flashed for 250 ms, with no intervening gray period. Five spatial frequencies (0.02, 0.04, 0.08, 0.16, 0.32 cycles per degree), four phases (0, 0.25, 0.5, 0.75), and six orientations (separated by 30°) were used. Each grating condition was presented approximately 50 times in random order.

The natural scenes stimulus consisted of 118 natural images taken from the Berkeley Segmentation Dataset (Martin et al., 2001), the van Hateren Natural Image Dataset (van Hateren and van der Schaaf, 1998), and the McGill Calibrated Colour Image Database (Olmos and Kingdom, 2004). The images were presented in grayscale and were contrast normalized and resized to 1174 × 918 pixels. The images were presented in a random order for 0.25 seconds each, with no intervening gray period.

Two natural movie clips were taken from the opening scene of the movie Touch of Evil (Welles, 1958). Natural Movie One was a 30 second clips repeated 20 or 30 times (2 or 3 blocks of 10), while Natural Movie Three was a 120 second clip repeated 10 times (2 blocks of 5). All clips were contrast normalized and were presented in grayscale at 30 fps.

### Spikes-to-Calcium Forward Model

All synthetic fluorescence traces were computed using MLSpike model number 3 (Deneux et al., 2016). This version models the supra-linear behavior of the calcium fluorescence response function in the most physiological manner (out of the three models compared) by (1) explicitly accounting for cooperative binding between calcium and the indicator via the Hill equation and (2) including an explicit rise time, *τ*_ON_

The model had seven free parameters: decay time (*τ*), unitary response amplitude (*A*), noise level (*σ*), Hill exponent (*n*), ΔF/F rise time (*τ*_ON_), saturation (*γ*), and baseline calcium concentration (*c*_0_). The last four parameters were fit on a ground truth dataset comprising 14 Emx1-Ai93 (from 9 individual neurons across 2 mice) and 17 Cux2-Ai93 recordings (from 11 individual neurons across 2 mice), each between 120 s and 310 s in duration, with simultaneous cell-attached electrophysiology and two-photon imaging (noise-matched to the ophys dataset) (Ledochowitsch et al., 2019): *n* = 2.42, *τ*_ON_ = 0.0034, *γ* = 0.0021, and *c*_0_ = 0.46. Reasonable values for the first three parameters were established by applying the MLSpike autocalibration function to all neurons recorded in the ophys dataset, computing a histogram for each parameter, and choosing the value corresponding to the peak of the histogram, which yielded *τ* = 0.359, *A* = 0.021, and *σ* = 0.047.

To convert spike times to synthetic fluorescence traces, MATLAB code publicly released by Deneux et al. (github.com/MLSpike/spikes) was wrapped into a Python (v3.6.7) module via the MATLAB Library Copiler SDK, and run in parallel on a high-performance compute cluster.

### 𝓁_0_-Regularized Event Extraction

Prior to computing response metrics, the normalized fluorescence traces for both the experimental and synthetic ophys data were passed through an 𝓁 _0_ event detection algorithm that identified the onset time and magnitude of transients (Jewell et al., 2018; Jewell and Witten, 2018; de Vries et al., 2020), using a revised version of this algorithm available at github.com/jewellsean/FastLZeroSpikeInference. The half-life of the transient decay was assumed to be fixed at 315 ms. To avoid overfitting small-amplitude false-positive events to noise in the fluorescence trace, the 𝓁 _0_-regularization was adjusted for each neuron such that the smallest detected events were at least 200% of the respective noise floor (computed as the robust standard deviation of the noise via the `noise std()` Python function from the `allensdk.brain observatory.dff` module) using an iterative algorithm. Note that this resulted in the artificial truncation of the event magnitude distribution for small event magnitudes evident in Figure 2B. All analysis was performed on these events, rather than continuous fluorescence time series.

### Visual Response Metrics

All response metrics were calculated from data stored in NWB 2.0 files (Ruebel et al., 2019) using Python code in a custom branch of the AllenSDK (github.com/jsiegle/AllenSDK/tree/ophys-ephys), which relies heavily on NumPy (van der Walt et al., 2011), SciPy (SciPy 1.0 Contributors et al., 2020), Matplotlib (Hunter, 2007), Pandas (McKinney, 2010), xarray (Hoyer and Hamman, 2017), and scikit-learn (Pedregosa et al., 2011) open-source libraries. For a given *ophys* stimulus presentation, the response magnitude for one neuron was defined as the summed amplitude of all of the events occurring between the beginning and end of the presentation. For a given *ephys* stimulus presentation, the response magnitude for one neuron was defined as the number of spikes occurring between the beginning and end of the presentation. Otherwise, the analysis code used for the two modalities was identical.

#### Responsiveness

To determine whether a neuron was responsive to a given stimulus type, the neuron’s response to its preferred condition was compared to a distribution of its activity during the nearest epoch of mean-luminance gray screen (the “spontaneous” interval). This distribution was assembled by randomly selecting 1000 intervals with the same duration of each presentation for that stimulus type (drifting gratings = 2 s, static gratings = 0.25 s, natural scenes = 0.25 s, natural movies = 1/30 s). The preferred condition is the stimulus condition (e.g., a drifting grating with a particular direction and temporal frequency) that elicited the largest mean response. The response reliability was defined as the percentage of preferred condition trials with a response magnitude larger than 95% of spontaneous intervals. A neuron was deemed responsive to a particular stimulus type if its response reliability was greater than 25%. Selectivity and preference metrics were only analyzed for responsive neurons.

#### Selectivity

The selectivity of a neuron’s responses within a stimulus type was measured using a lifetime sparseness metric (Vinje and Gallant, 2000). Lifetime sparseness is defined as:

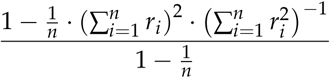

where *n* is the total number of conditions, and *r*_*i*_ represents the response magnitude for condition *i*. If a neuron has a non-zero response to only one condition (maximally selective response), its lifetime sparseness will be 1. If a neuron responds equally to all conditions (no selectivity), its lifetime sparseness will be 0. Importantly, lifetime sparseness is a nonparametric statistic that considers a neuron’s selectivity across all possible stimulus conditions within a stimulus type, rather than conditions that vary only one parameter (e.g., orientation selectivity). For that reason, it is applicable to any stimulus type.

#### Preference

For all stimulus types, the preferred condition was defined as the condition (or frame, in the case of natural movies) that elicited the largest mean response across all presentations. For drifting gratings, the preferred temporal frequency was defined as the temporal frequency that elicited the largest mean response (averaged across directions). For static gratings, the preferred spatial frequency was defined as the spatial frequency that elicited the largest mean response (averaged across orientations and phases).

### Matching Layer Distributions

Neurons in the ophys dataset were assigned to layers based on the depth of the imaging plane (<200 µm = L2/3, 200-325 µm = L4, 325-500 µm = L5, >500 µm = L6), or the mouse Cre line (Nr5a1-Cre and Scnn1a-Tg3-Cre neurons were always considered to be L4). Neurons in the ephys dataset were assigned to layers after mapping their position to the Common Coordinate Framework version 3 (Wang et al., 2020). CCFv3 coordinates were used as indices into the template volume in order to extract layer labels for each cortical unit (see Siegle et al., 2019 for details of the mapping procedure).

To test for an effect of laminar sampling bias, L6 neurons were first removed from both datasets. Next, since the ephys dataset always had the highest fraction of neurons L5, neurons from L2/3 and L4 of the ophys dataset were randomly sub-sampled to match the relative fraction of ephys neurons from those layers. The final resampled layer distributions are shown in Figure S1B.

### Burst Metrics

Bursts were detected using the *LogISI* method (Pasquale et al., 2010). Peaks in the histogram of the log-adjusted inter-spike intervals (ISI) were identified, and the largest peak corresponding to an ISI of less than 50 ms was set as the intra-burst peak. In the absence of such a peak, no bursts were found. Minima between intra-burst peak and subsequent peaks were found, and a void parameter, representing peak separability, was calculated for each minimum. The ISI value for the first minimum where the void parameter exceeds a default threshold of 0.7 was used as the *maxISI*-cutoff for burst detection. Bursts were then defined as a series of >3 spikes with ISIs less than *maxISI*. If no cutoff was found, or if *maxISI* > 50 ms, burst cores were found with <50-ms ISI, and any spikes within maxISI of burst edges were included.

R code provided with a comparative review of bursting methods (Cotterill et al., 2016) (github.com/ellesec/burstanalysis) was wrapped into Python (v.3.6.7) using the rpy2 interface (rpy2.github.io), and run in parallel on a high-performance compute cluster.

### Statistical Comparisons

Jensen-Shannon distance was used to quantify the disparity between the distributions of metrics from ophys and ephys. This is the square root of the Jensen-Shannon divergence, also known as the total divergence to the mean, which, while derived from the Kullback–Leibler divergence, has the advantage of being symmetric and always has a finite value. The Jensen-Shannon distance constitutes a true mathematical metric that satisfies the triangle inequality. (Lin, 1991). We used the implementation from the SciPy library (SciPy 1.0 Contributors et al., 2020) (`scipy.spatial.jensenshannon`). For selectivity and responsiveness metrics, Jensen-Shannon distance was calculated between histograms with 10 equal-sized bins between 0 and 1. For preference metrics, Jensen-Shannon distance was calculated between the preferred condition histograms, with unit spacing between the conditions. To compute *P* values for Jensen-Shannon distances, we used a bootstrap procedure to randomly sub-sample metric values from one modality, and calculated the probability that the true inter-modality distance would be less than the distance between the distributions of two non-overlapping intra-modality samples.

The Pearson correlation coefficient (`scipy.stats.pearsonr`) was used to quantify the correlation between two variables. The Mann–Whitney *U* test (`scipy.stats.ranksums`) was used to test for differences in running speed or running fraction between the ophys and ephys datasets.

### Clustering of Response Reliabilities

We performed a clustering analysis using the response reliabilities by stimulus for each neuron (defined as the percentage of significant trials to the neuron’s preferred stimulus condition), across drifting gratings, static gratings, natural scenes, and natural movies. We combined the reliabilities for natural movies by taking the maximum reliability over Natural Movie One and Natural Movie Three. This resulted in a set of four reliabilities for each neuron (for drifting gratings, static gratings, natural movies, and natural scenes).

We performed a Gaussian Mixture Model clustering on these reliabilities for cluster numbers from 1 to 50, using the average Bayesian Information Criterion (Schwarz, 1978) on held-out data with four-fold cross validation to select the optimal number of clusters. Once the optimal model was selected, we labeled each cluster according to its profile of responsiveness (i.e. the average reliability across all neurons in the cluster to drifting gratings, static gratings, etc.), defining these profiles as “classes”. For each neuron, we predicted its cluster membership using the optimal model, and then the class membership using a predefined responsiveness threshold. We repeated this process 100 times to estimate the robustness of the clustering and derive uncertainties for the number of cells belonging to each class.

## Data and Code Availability

Code and intermediate data files required to generate all manuscript figures will be made available through the following repository: https://github.com/AllenInstitute/ophys_ephys_comparison_paper

NWB files for the ophys and ephys sessions (excluding the GCaMP6f ephys experiments) are available via the AllenSDK (https://github.com/AllenInstitute/AllenSDK), and also as an Amazon Web Services Public Data Set (https://registry.opendata.aws/allen-brain-observatory/).

NWB files for the GCaMP6 ephys experiments will be released prior to publication.

## Supplementary Figures

**Figure S1:**
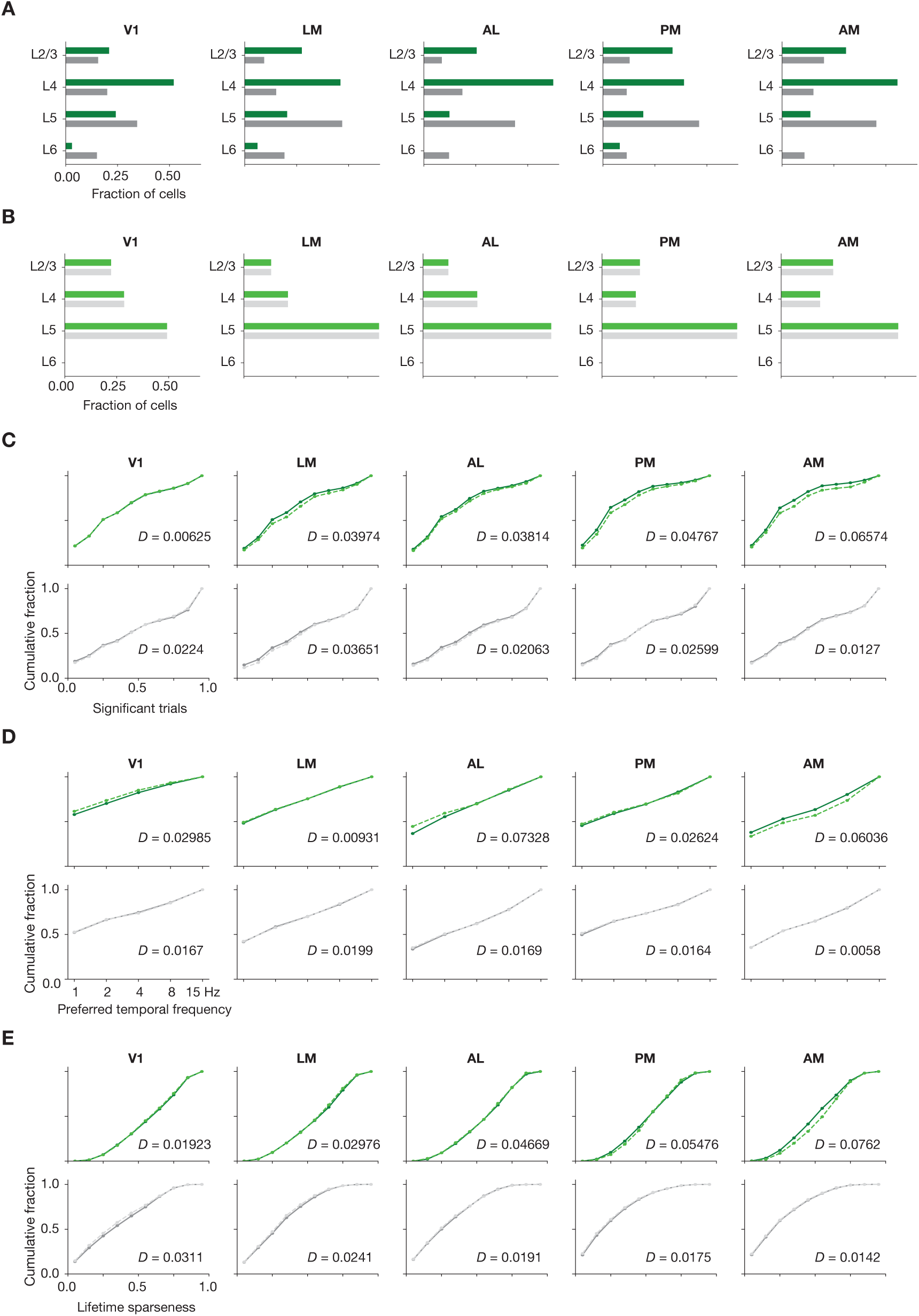
Matching laminar distribution patterns across modalities. (A) Original layer distributions, prior to resampling, for ophys (green) and ephys (gray). (B) Layer distributions following the resampling procedure, for ophys (light green) and ephys (light gray). (C) Cumulative histograms of significant trial fractions for ophys and ephys, before and after resampling. (D) Cumulative histograms of preferred temporal frequency for ophys and ephys, before and after resampling. (E) Cumulative histograms of preferred temporal frequency for ophys and ephys, before and after resampling. The value D represents the Jensen-Shannon distance between the before/after distributions. In panels (C-E), lighter dashed lines represent the values after resampling.

**Figure S2:**
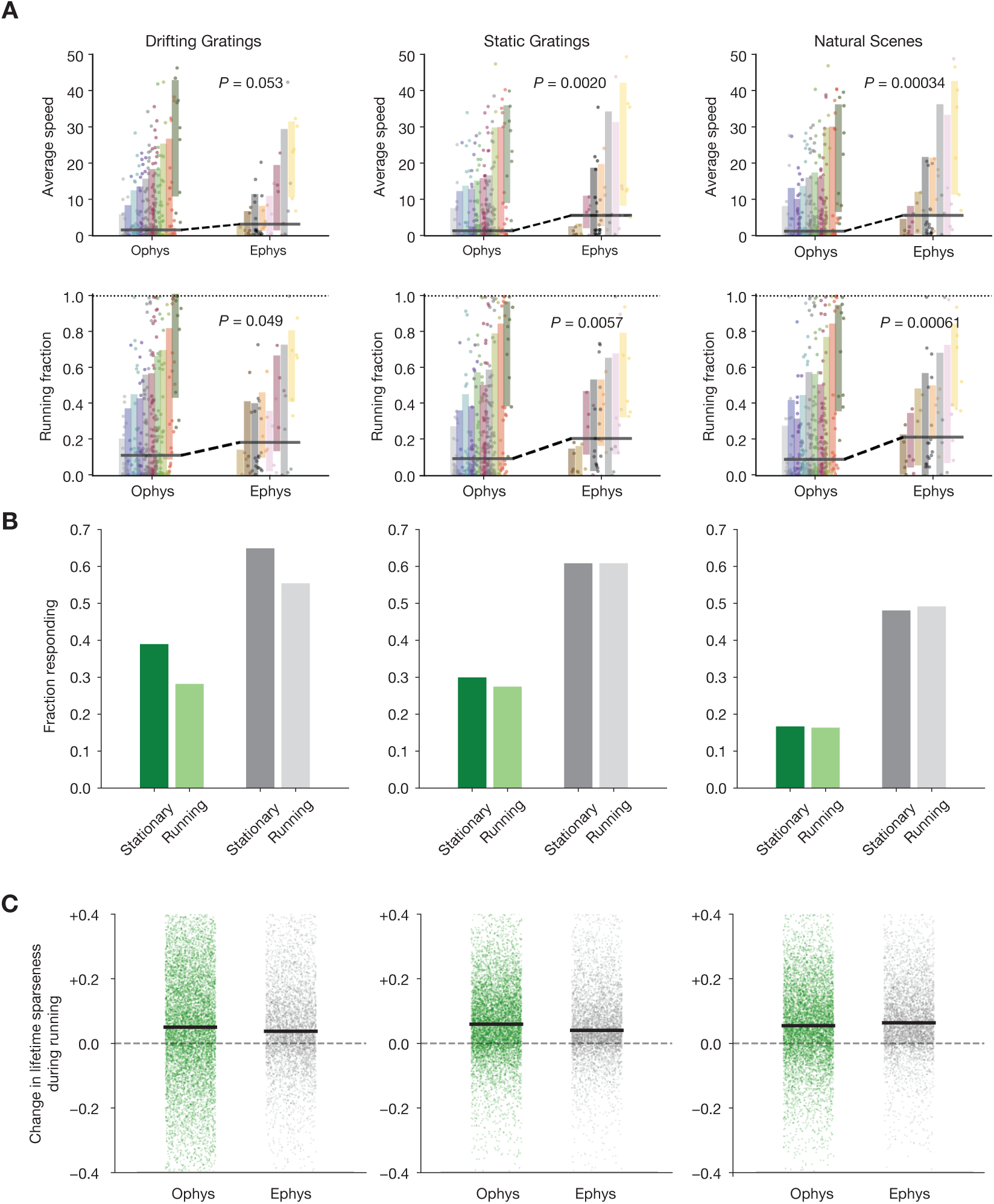
Characterizing the impact of running behavior on response metrics. (A) Average speed and running fraction for individual mice from ophys and ephys experiments, grouped by genotype. Shaded areas represent the mean ± standard deviation for each genotype. Colors are the same as in Figure 1F. P values are from the Mann–Whitney U test between all ophys and all ephys values for each stimulus/metric combination. (B) Fraction of neurons responding to three stimulus types during stationary and running intervals. Neurons were included only if the mouse was stationary or running on at least 20% of trials. (C) Change in lifetime sparseness during running intervals for all neurons from ophys and ephys experiments. Black line represents the median change for each modality/stimulus type.

**Figure S3:**
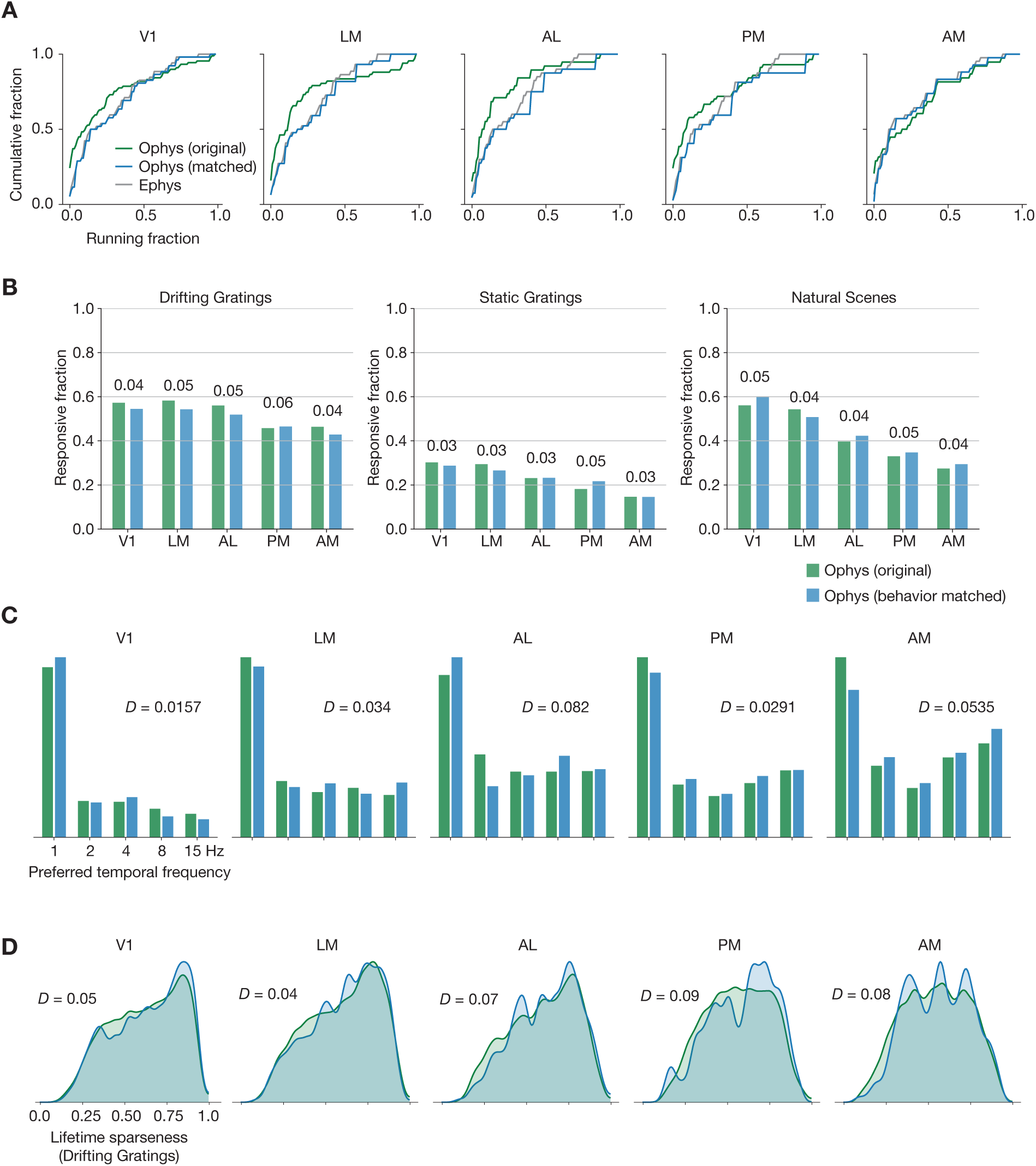
Matching running behavior across modalities. (A) Cumulative histogram of drifting gratings running fractions for the ephys dataset (gray), and for the ophys dataset before (green) and after (blue) performing the behavioral matching procedure. (B) Fraction of neurons deemed responsive to each of three stimulus types, before and after sub-sampling the ophys experiments to match ephys running behavior. Numbers above each pair of bars represent the Jensen-Shannon distance between the full distribution of responsive trial fractions for each stimulus/area combination. (C) Distribution of preferred temporal frequencies for all neurons in 5 different areas. The Jensen-Shannon distance between the original ophys and resampled ophys distributions is shown for each plot. (D) Distributions of lifetime sparseness in response to a drifting grating stimulus for all neurons in 5 different areas. The Jensen-Shannon distance between the original ophys and resampled ophys distributions is shown for each plot.

**Figure S4:**
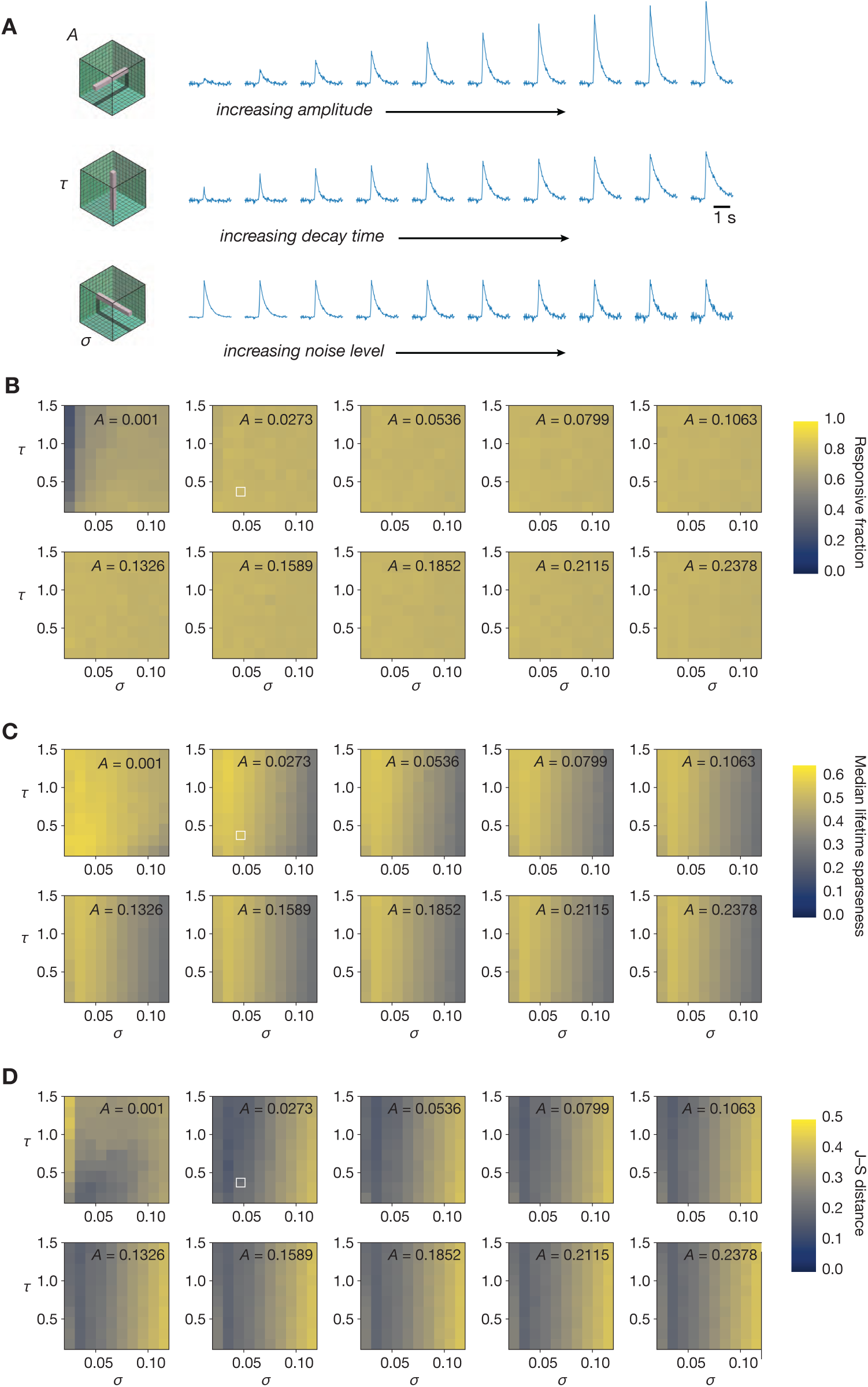
Sampling the entire space of forward model parameters. (A) Schematic of three different “slices” through forward model parameter space, sampling all values of unitary amplitude (*A*), decay time (*τ*), or noise level (*σ*). (B) Fraction of neurons responding to drifting gratings for all parameter combinations. (C) Median lifetime sparseness in response to a drifting grating stimulus for all parameter combinations. The median value for the experimental ophys data determines the maximum value of the color scale. (D) Jensen-Shannon distance between the distributions of drifting gratings lifetime sparseness for all parameter combinations. In panels (B–D), the parameter combination used in Figure 4 is highlighted in white.

**Figure S5:**
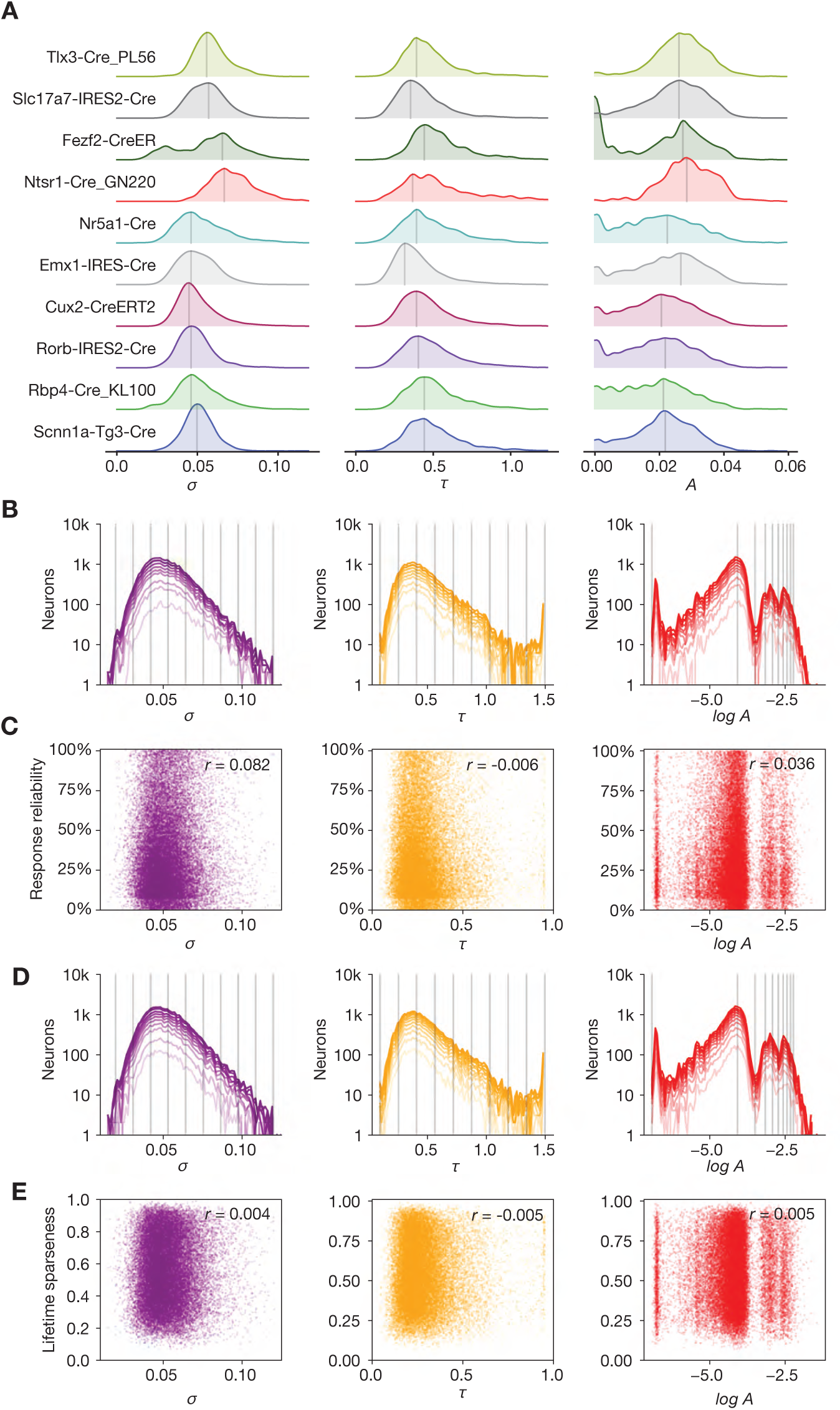
Relationship between forward model parameters, genotype, and functional metrics. (A) Distribution of noise level (*σ*), decay time (*τ*), and amplitude (*A*) parameters for the 10 different Cre lines used in this study. (B) Effect of sub-selecting neurons based on drifting gratings response reliability. Each line represents the parameter distribution for quantiles of increasing reliability (from dark to light). (C) Relationship between forward model parameters and response reliability. (D) Effect of sub-selecting neurons based on selectivity. Each line represents the parameter distribution for quantiles of increasing selectivity (from dark to light). (E) Relationship between forward model parameters and drifting gratings lifetime sparseness.

